# Perturbational fitness analysis of CRISPR screens uncovers information-theoretic relation between gene function and selection

**DOI:** 10.1101/2025.04.29.651199

**Authors:** A.N. Andersen, N. Chica, L. Piechaczyk, S. Nakken, M. Zucknick, J.M. Enserink

**Affiliations:** Department of Molecular Cell Biology, Institute for Cancer Research, Oslo University Hospital, 0379 Oslo, Norway; Centre for Cancer Cell Reprogramming, Institute of Clinical Medicine, Faculty of Medicine, University of Oslo, 0318 Oslo, Norway; Section for Biochemistry and Molecular Biology, Faculty of Mathematics and Natural Sciences, University of Oslo, 0371 Oslo, Norway; Department of Tumor Biology, Institute for Cancer Research, Oslo University Hospital, 0379 Oslo, Norway; Centre for Bioinformatics, Department of Informatics, University of Oslo, Oslo, Norway; Oslo Centre for Biostatistics and Epidemiology, University of Oslo, 0317 Oslo, Norway; Faculty of Medicine, University of Oslo, 0318 Oslo, Norway

## Abstract

Despite major advances in genetic screening technology, a formal approach for quantifying gene function remains underdeveloped, thereby limiting the utility of these techniques in deciphering the complex behavior of human cells. In this study, we leverage information theory with a perturbational analysis of replicator dynamics to characterize functional drivers of selection in pooled CRISPR screens. Our approach challenges established methods for CRISPR screen analysis, while offering additional insight into selection dynamics through the Kullback-Leibler divergence (*D*_*KL*_) and cumulants of the fitness distribution. By modeling fluctuations in gene-fitness effects as a linear response to environmental perturbations, we derive a geometric measure for genomic information content based on a second-order approximation of the *D*_*KL*_. Our analysis reveals that functional information—encoded (or shared) between genes—can be quantified by analyzing the directions corresponding to maximal conditional selection within the space of decomposed gene-environment interactions. This geometric representation offers several advantages for the functional analysis of the human genome and its network architecture. Moreover, by constraining the space to cell-type-specific fluctuations, we uncover developmental and tissue-specific functional signatures. These findings represent significant progress in the dynamic analysis of gene function and in the functional wiring of the human genome.

## I. INTRODUCTION

Functional reverse engineering of the human genome remains a major challenge in modern biological science due to the vastness and complexity of genetic relations that contribute to phenotypic variability in human cells. Despite significant advances in high-throughput genetic screening, current analytical tools are largely provisional, and a standardized language for quantifying gene functions has yet to be established [1–3].

Modern evolutionary theory links the selection of complex phenotypes to the fitness currency of individual genotypes, which contribute to growth under various environmental constraints [4, 5]. By extension, the function of genes can be understood from the set of gene-environment interactions shaping cellular or organismal fitness. This principle has been applied to deconvolve functional similarities from genetic perturbation screens by measuring correlations in gene–fitness associations across numerous cell culture conditions or genetic environments [6–8]. Recently, pooled CRISPR screening in human cancer cells has become a crucial approach to systematically measuring genetic dependencies and exploring functional similarities [9–11].

CRISPR screens track the selection of sgRNA-targeted, single-gene deletion mutants over time by quantifying the relative abundance of each sgRNA in a population through next-generation sequencing while assuming proportionality between cell representation and sgRNA frequency. Consequently, in principle, CRISPR screens can be analyzed using statistical models of natural selection, mapping frequency changes to growth rates, phenotype-fitness covariances, and information gain in response to selection. However, the most widely used methods typically compute a fitness or essentiality score with varying degrees of formality, while controlling for confounders such as variations in sgRNA efficacy, copy number effects, sequence read distributions, sampling time, and batch issues [12–17]. With the detection of core essential genes being the primary benchmark for assessing these methods and with limited focus on a principled modeling of selection dynamics, less is known about the preservation of information gained from variations in selection pressure or conditional changes in fitness distributions.

Research on retrieving gene functional information has focused on post-processing fitness scores in ensembles of screened cancer cells before computing linear correlations between genes. CRISPR screen data are fraught with technical covariances, which confound computed gene correlations and bias the detection of functional subgroups, thereby masking other valuable information [18–21]. Two main classes of strategies have been developed to address this challenge. One approach preserves average fitness effects per gene while removing the leading principal components dominating the covariance structure before reconstructing the data and analyzing correlations between genes [19, 22]. The other strategy involves computing covariance-normalized gene similarities to counter confounding cell line correlations, thereby enabling improved enrichment of functional relations and genome-wide mapping of cellular functions [20]. This latter approach is a form of whitening transformation in which similarities are probed from the latent structure of independent gene dependencies [23, 24]. However, relying strictly on similarity coefficients between genes may limit progress in understanding how genetic variation in functional signatures leads to different phenotypes or how conditional rewiring of functions shapes the fitness landscape in different cellular contexts [25].

The diversity of characteristics needed to describe the consequences of different genetic alterations poses a central challenge to the development of a universal system for gene functions [1]. Any particular measure will prioritize genes based on their functional proximity to a phenotype, such as essential genes, when evaluating perturbations in growth. Building on recent advances that connect selection principles with information theory, we employ the information encoded in genes as a unitless metric of functional relevance in a given environment [26–30]. By using a perturbational formulation of the replicator equation to analyze CRISPR screens, we show how information gain—captured by the *D*_*KL*_ and its second-order approximation—can be used to characterize experimental selection pressure and bounds on functional genomic fitness associations. This framework is simpler than alternative methods and, with the successful layering of techniques to counter confounding effects, facilitates a direct interpretation of the computed metrics. Furthermore, by considering selection as a process that reduces uncertainty about environmental fluctuations (quantified through mutual information), we pinpoint gene function signatures as directions in fitness component space, where the maximum information encoded in every gene corresponds to the squared distance of their normalized projection coordinates. We proceed to examine these signatures with respect to pairwise gene relations, the limits of functional enrichment, functional network structure, and cell-type conditional networks.

## II. RESULTS

### A. Simple fitness perturbation analysis (FPA)

Pooled CRISPR screens estimate fitness perturbations in response to sgRNA-targeted gene deletions by tracking changes in the relative frequency of sgRNA sequence reads in a mixed population of mutant cells [31]. Thus, under typical cell culture conditions, where cells can grow exponentially with limited density constraints, the frequency dynamics of mutants can be described by the simple replicator equation 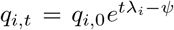. Here, *λ*_*i*_ is the growth rate of mutant *i* in a genome-wide population of *N* distinct mutants (*i* ∈ *G*), and 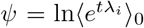 is the associated cumulant generating function.

The expected population dynamics can be described by expanding *ψ*, defined as 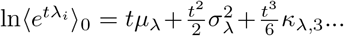 where *µ*_*λ*_ is the expected growth rate, 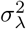 is the variance, *κ*_*λ*,3_ is the skewness, etc., evaluated at *t* = 0. The effects of these parameters on changes in mutant frequencies, or overall selection, can be summarized using the *D*_*KL*_,

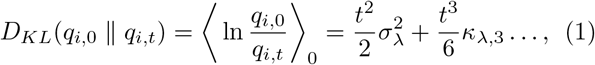

relating the expected divergence to the variation in growth rates. See Methods section C for extensions to related divergence metrics and Fisher’s fundamental theorem. This derivation offers a useful formalism for characterizing CRISPR screen selection. If the second-order approximation is valid (i.e.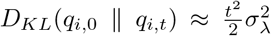, then the fitness distribution is effectively normal, implying more drift and less selective pressure described by higher-order cumulants. Conversely, in environments where certain gene functions are selectively important, the tails of the fitness distribution become more influential over time, violating the second-order approximation. This solution also provides a straightforward equality between perturbations in growth rate and mutant frequency dynamics:

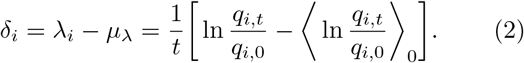

Here, *δ*_*i*_ is the slope of the log-fold change in frequencies adjusted by the population average, quantifying the fitness effect of gene *i*. Intuitively, if *δ*_*i*_ = 0, gene *i* exerts no functional effect in the given environment.

The above framework can be extended to conditional fitness perturbations, with 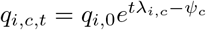 and *λ*_*i,c*_ = *f*_*i,c*_ + *λ*_*i*_. Here, *f*_*i,c*_ represents a gene-environment interaction, and its perturbation from the population average can be calculated as the differential slope of selection in condition *c* relative to a reference control: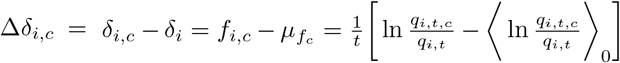. In this case, Δ*δ*_*i,c*_ = 0 implies that gene *i* has no unique functional contribution to the cellular response to *c*, whereas positive or negative values represent a genetic interaction effect [7].

By interpreting *q*_*i,t,c*_ and *q*_*i,t*_ as conditional and marginal probabilities, respectively, ln 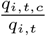 is the *pointwise specific information I*_*t*_(*i*; *C* = *c*) [32, 33], and Δ*δ*_*i,c*_ measures the non-redundant information that mutation *i* provides about *c* (see Methods section D). Moreover, if the baseline fitness and the interaction effects are independent, the *specific information* gained by a population under selection *c* can be approximated variationally: *I*_*t*_(*G*; *C* = *c*) = *D*_*KL*_(*q*_*i,c,t*_ ∥ *q*_*i,t*_) ≈ *D*_*KL*_(*q*_*i*,0_ ∥ *q*_*i,c,t*_) −*D*_*KL*_(*q*_*i*,0_ ∥ *q*_*i,t*_). Thus, for pan-cancer screens in which a well-defined control condition does not exist, *D*_*KL*_(*q*_*i*,0_ ∥ *q*_*i,c,t*_) provides an empirically accessible metric to compare selective pressure and information gain across environments.

### B. D_KL_ and fitness distribution cumulants are important factors for assessing CRISPR screen characteristics and hit sensitivity

To gain insight into the utility of the outlined fitness perturbation analysis (FPA), we evaluated this framework by applying it to controlled stochastic simulations, time-resolved CRISPR screen experiments performed under defined treatment conditions, and large-scale pancancer CRISPR screens (Fig. 1).

**Fig. 1.**
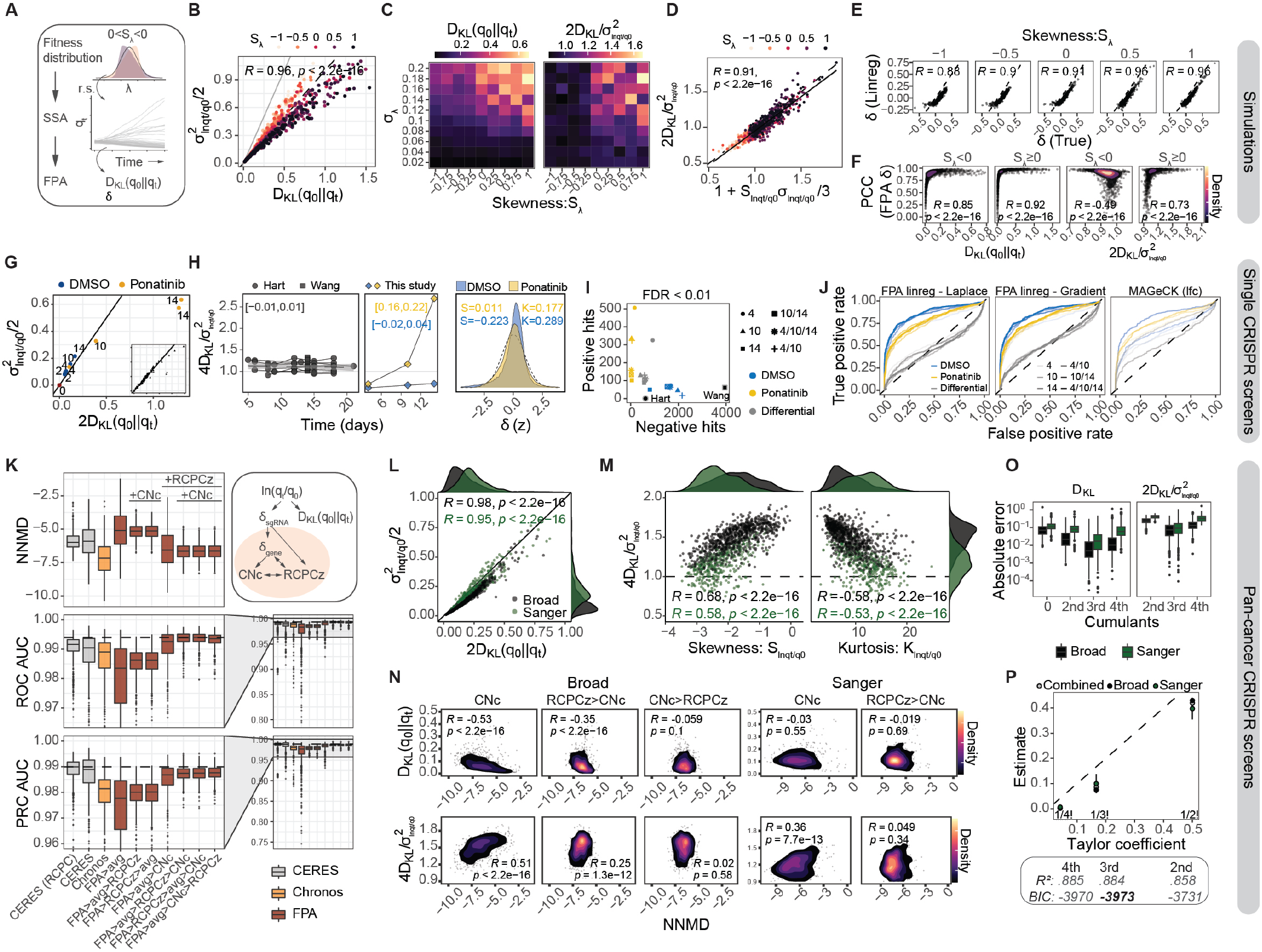
Perturbational analysis of CRISPR screen selection dynamics. **A** Overview of the simulation experiments (panels A–F, Fig. S1). **B** *D*_*KL*_ versus the empirical variance for different fitness skewness thresholds; *R* indicates the Pearson correlation. **C** Average *D*_*KL*_ (*left*) and relative divergence (*right*) for various fitness variance and skewness thresholds. **D** Relative divergence estimated from the empirical skewness; *R* indicates the Pearson correlation. **E** Comparison between regression-estimated and true fitness across different fitness skewness thresholds; *R* indicates the Pearson correlation. **F** Pearson correlations for FPA-estimated fitness with respect to *D*_*KL*_ and relative divergence under positive and negative fitness skewness; *R* indicates the Spearman correlation. **G** *D*_*KL*_ plotted against the empirical variance for CRISPR screen samples under selective pressure (Ponatinib) or left untreated (DMSO). The inset includes all single-condition CRISPR screen experiments from Fig. S2B. **H** (*Left*) Relative divergence as a function of time (values in brackets indicate the 95% CI for the slope). (*Right*) Standardized fitness distributions at day 14, with values indicating skewness (S) and kurtosis (K). The dashed line represents a standard normal distribution. **I** Shift in the number of positive and negative significant perturbations under selective pressure. Black dots indicate the average from other CRISPR screens. **J** ROC curves for detecting essential genes using FPA with different smoothing techniques and mean as a gene-level statistic, or MAGeCK (log-fold changes). **K** NNMD, ROC, and PRC AUC for detecting essential genes using fitness scores computed via CERES, Chronos, or FPA on the Broad Institute dataset. The *upper right* panel outlines the workflow for FPA combined with CNc and RCPCz. **L** *D*_*KL*_ with respect to the empirical variance, and **M** relative divergence with respect to the empirical skewness and kurtosis for both the Broad and Sanger Institute datasets; *R* indicates Pearson correlation. **N** NNMD for detecting essential genes with respect to *D*_*KL*_ or relative divergence; *R* indicates Spearman correlation. **O** Absolute error in expressing *D*_*KL*_ (*left*) or relative divergence (*right*) as a function of the number of cumulants. **P** Estimated Taylor coefficients from regressing *D*_*KL*_ using the 2nd–4th cumulants. The box shows *R*^2^ and *BIC* for regression fits that use an increasing number of cumulants.

#### 1. Stochastic simulation experiments

Simulation experiments were performed using the stochastic simulation algorithm [34], where populations of replicators were sampled from predefined fitness distributions with varying mean, variance, and skewness (Fig. 1A) while controlling for sources of technical noise and bias. Comparing the overall selection, given by *D*_*KL*_(*q*_*i*,0_ ∥ *q*_*i,t*_), to an empirical estimator of the second cumulant, 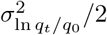 (see Methods section B), revealed a consistent linear correspondence (Fig. 1B), with deviations determined by fitness skewness (Fig. S1A-D).Consistent with the theory, fitness variance and positive skewness synergized to increase overall selection (Fig. 1C, *left*). Meanwhile, a relative divergence, 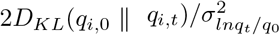, isolated higher-order selective forces and was influenced only by the fitness skewness (Fig. 1C, *right*), irrespective of sampling time (Fig. S1C-D). Interestingly, increasing the average growth rate, shifts in the T0 reference point, and sampling bottlenecks each reduced relative divergence (Fig. S1B-E), except for multiple consecutive bottlenecks, which produced the opposite effect (Fig. S1B). Finally, empirical estimation of the third cumulant explained the simulated variation in relative divergence (Fig. 1D and S1L), further supporting the outlined framework for analyzing selection dynamics. To infer perturbations in growth rates, we used Equation (2) directly for single time points and developed two methods for time series data: one estimates the slope via linear regression, and the other uses singular value decomposition (SVD; see Methods). Both methods produced correlated estimates of true fitness and only diverged slightly with changes in skewness; linear regression was more resistant to noise (Fig. 1D and S1F-K). Unsurprisingly, overall selection driven by the underlying fitness variance strongly determined the extent to which fitness perturbations could be estimated (Fig. 1E and S1F). In contrast, the accuracy of the fitness estimation decreased under conditions where the relative divergence converged toward the second-order approximation 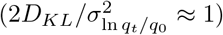.

#### 2. Single CRISPR screens

To evaluate this framework for analyzing the selection dynamics of real CRISPR screens, we retrieved publicly available high-quality data from two studies by Hart et al. and Wang et al. [9, 10]. In addition, we performed a controlled, time-resolved CRISPR screen to examine the impact of selective pressure on a well-characterized driver of cell proliferation. Specifically, we stably transformed Ba/F3 cells to overexpress the human BCR-ABL1-T315I oncogene, which encodes a constitutively active tyrosine kinase that induces oncogene addiction in these cells [35]. Using a genome-wide sgRNA library, we generated a population of mutant cells and treated them with DMSO (carrier control) or sublethal concentrations of Ponatinib, a tyrosine kinase inhibitor used clinically to treat T315I-mutant chronic myeloid leukemia (CML).

Remarkably, across all CRISPR screens, *D*_*KL*_ scaled with the second cumulant almost precisely by a factor of 1*/*2 (Fig. 1G and S2A-D), reflecting a negative fitness skewness for most conditions. Moreover, applying selective pressure with Ponatinib increased both overall selection and relative divergence in BCR-ABL1-T315I-transformed Ba/F3 cell populations. Whereas untreated screens showed constant selective pressure, Ponatinib treatment increased relative divergence over time, consistent with a shift from negative to positive skewness and a rise in the proportion of significantly positive fitness perturbations (Fig. 1H-I and S2E-F).

The selective pressure from Ponatinib treatment reduced the sensitivity to detect essential genes (Fig. 1J and S2G-H). This suggests that, despite increased variance, the distributional shift in fitness made it difficult to separate constitutively core essential (CCE) genes from non-essential (NES) genes. To ensure this reduction was not caused by a treatment-specific bias in either group, we also performed the same tests on conditional fitness perturbations, which revealed no systematic separation between the groups (Fig. 1H).

Because zero read counts are common, a +1 pseudocount (Laplace smoothing) is typically used but can cause time-dependent shifts due to selection; therefore, we tested a gradient smoothing method that reduced these shifts and slightly improved CCE detection under Ponatinib (Fig. 1J, S2A–D, S2G–H). Surprisingly, the widely used hit-calling algorithm MAGeCK [12] underperformed both FPA methods to detect CCEs (Fig. 1J, S2G–I) and showed highly time-varying performance. These findings indicate that FPA provides a more robust analysis overall and that essential gene detection depends on selective pressure, which can be revealed using the *D*_*KL*_ and its relative divergence from the fitness variance.

#### 3. Pan-cancer CRISPR screens

To assess the use and interpretation of *D*_*KL*_ in heterogeneous selection environments, we analyzed two large pan-cancer datasets from the Broad and Sanger Institutes for the Cancer Dependency Map [15, 36]. For this analysis, we also considered that fitness perturbations would be heavily confounded by gene copy numbers (CNs) and batch effects. Batch effects, in particular, can be effectively mitigated by removing the confounding principal components (RCPC) [19]. We applied this RCPC procedure and additionally rescaled the data using gene-wise fitness variances (RCPCz; Fig. S3A). For CN correction (CNc), we performed a linear mixed-model regression with random intercepts and slopes, grouping at the gene level to account for genomic variation in cutting toxicity [37]. We then reconstructed the data, excluding both the mean and gene-wise CN effects (Fig. S3A).

Compared with state-of-the-art methods such as CERES and Chronos [15, 17], FPA detected essential genes with similar precision and sensitivity. Moreover, when combined with RCPCz, FPA showed a much lower proportion of weakly performing screens (Fig. 1K). FPA with CNc also reduced the correlations between CNs and gene-level fitness effects, similar to what has been reported for Chronos, yet did not incur the same drop in precision–recall (Fig. S3C and 1K) [17]. Interestingly, the effect of these correction modules appeared to be order-invariant and could seamlessly replace less accurate methods (Fig. S3D) [38].

For these datasets, *D*_*KL*_ also scaled with the second cumulant by a factor of 1*/*2, while the Broad dataset exhibited lower overall selection and higher relative divergence (Fig. 1L). For both datasets, the relative divergence was positively correlated with skewness and negatively correlated with the kurtosis (Fig. 1M). Consistent with previous observations, these changes were accompanied by a reduction in essential gene detection sensitivity as relative divergence increased (Fig. 1N and S3E-F), an effect mitigated by RCPCz.

The variation in *D*_*KL*_ was largely explained by the second and third cumulants. Skewness accounted for most of the higher-order variation in selection pressure (Fig. 1O), and kurtosis aligned closely with its theoretical minimum as given by the Cauchy–Schwarz inequality (Fig. S4A-B) [39]. Moreover, estimating the Taylor coefficients for the 2nd–4th cumulants revealed proportionality with their theoretical values, but also highlighted a slight negative bias in the *D*_*KL*_, possibly related to technical influences or unaccounted higher-order effects (Fig. 1P and S1). Finally, experimental data matched with theoretical predictions comparing related divergence measures (Fig. S4C-F; see Methods sections C-D). Together, these data support the idea that CRISPR screens can be analyzed using an information-theoretic framework for selection.

### C. Genome-wide functional fitness associations relate to selection dynamics and information gain

To assess the link between genome-wide functional associations and observed selection dynamics, we performed directional gene ontology (GO) enrichment [40] on estimated fitness perturbations (*δ*) and interaction effects (Δ*δ*). We then quantified the proportion of significant GO terms (*p ≤* 0.01) among all tests (Fig. S5). Across all fitness metrics, screens with higher *D*_*KL*_ displayed more functional associations for both positive and negative fitness perturbations, whereas those with higher relative divergence had fewer (Fig. 2A, *left* and *middle* panels, S6A). This tendency was variably affected by RCPCz (Fig. S6A-B), which also led to some homogenization of cell-type-specific GO enrichment (Fig. S5C). However, upon scaling the *δ*-values by the original fitness variance (*σ*_*λ*_), the relative divergence revealed a shift from negative to positive functional associations for interaction effects (Δ*δ*) (Fig. 2A, *right* panels, S6B). This finding suggests that the selective pressure for a given cell type may be determined by the distribution of condition-specific sensitizers throughout the genome.

**Fig. 2.**
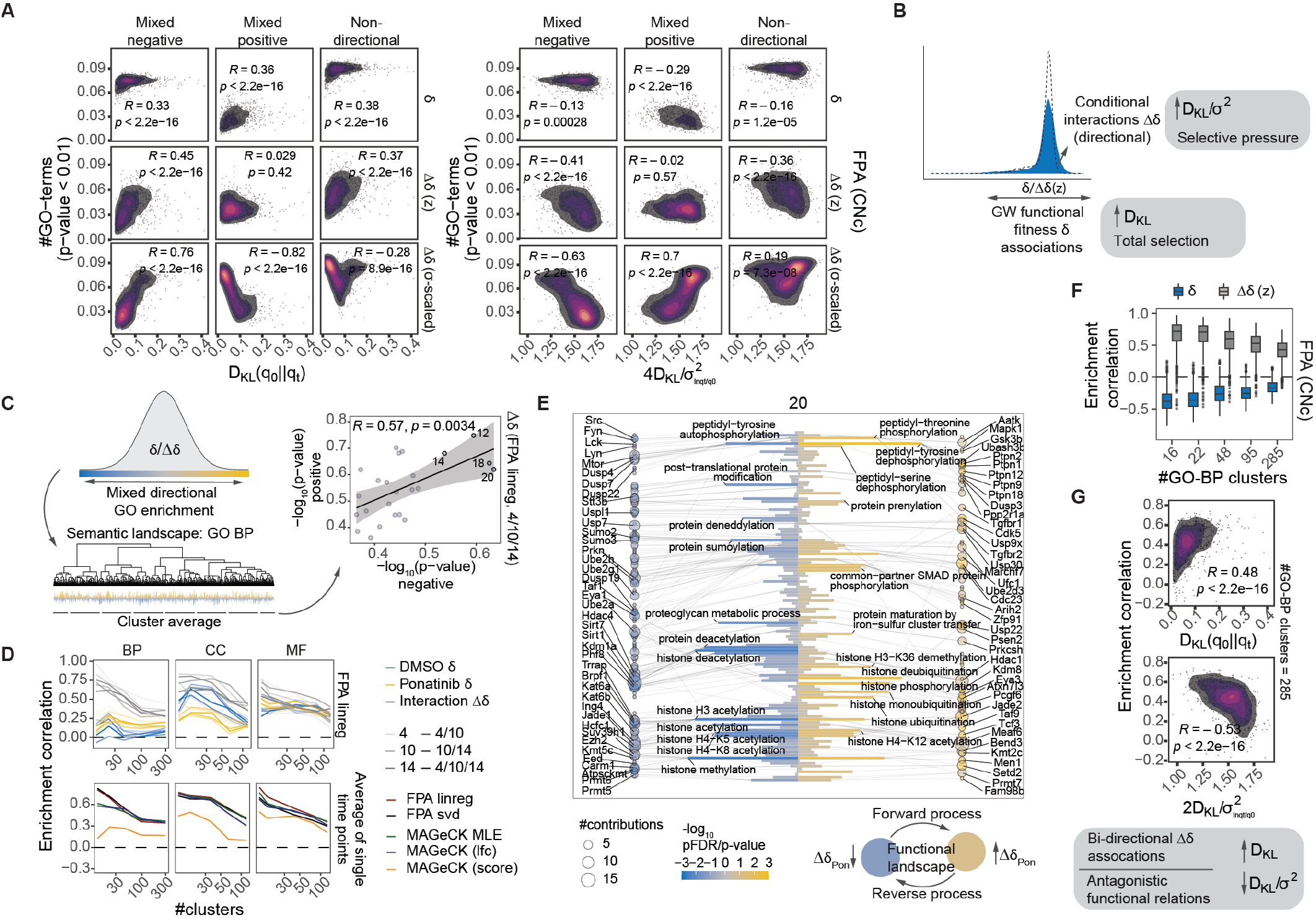
Functional associations with the D_KL_. **A** Fitness GO enrichment sensitivity with respect to *D*_*KL*_ and relative divergence; *R* indicates the Spearman correlation. **B** Summary of results from panel A and Fig. S6. *D*_*KL*_ is associated with stronger overall functional associations, while relative divergence quantifies a directional asymmetry in functional associations involving interaction effects. **C** Overview of the workflow for detecting bidirectional GO enrichment via semantic similarity clustering of GO terms (*left*). The average directional enrichment statistic is shown per semantic cluster; *R* indicates the Pearson correlation. **D** Directional enrichment correlations for various FPA-estimated fitness perturbations at different clustering depths (*top panel*), and comparison against MAGeCK-estimated differential test scores (*bottom panel*). **E** Directional GO enrichment results for cluster 20, indicating contributing genes. Antagonistic functions relevant to the condition appear diametrically associated with the interaction effects Δ*δ* (*bottom right*). **F** Directional enrichment correlations in the Broad Institute dataset across different clustering depths, **G** and with respect to *D*_*KL*_ and relative divergence; *R* indicates the Spearman correlation. Asymmetrical functional interaction effects correlate with higher relative divergence (*bottom*).

A hypothesis for the interaction effects is that they provide directional information about the functions that drive conditional selection. Although this idea has been extensively studied for genetic interactions in lower eukaryotes [6, 41], it is difficult to verify in complex cancer cell environments with poorly defined conditions. However, if correct, we should expect a bidirectional fitness association of antagonistic regulators belonging to the same cellular process. We tested this by grouping GO terms using semantic similarity [42] and computing correlations in directional enrichment (− log_10_(*p-value*)) across semantic clusters (Fig. 2C).

Using differential selection with Ponatinib as a reference, we found that only the interaction effects produced a substantial bidirectional enrichment of antagonistic cell processes with functional relevance to the treatment (Fig. 2D-E and S7A). For instance, deletion of tyrosine phosphatases that cause desensitization clustered together with several non-receptor tyrosine kinases that confer sensitization to the drug (Fig. 2E) [43]. However, this pattern also extended to several other clusters, including regulators of histone modification (Fig. S7C) [44]. This bidirectional enrichment was also evident for cancer-specific interaction effects, showing a positive association with the *D*_*KL*_ and a negative association with the relative divergence (Fig. 2F-G). These associations persisted even after RCPCz (Fig. S8). These results indicate that the *D*_*KL*_ and the relative divergence collectively capture the extent of genome-wide functional associations and the symmetry between cancer-specific suppressors and activators. Moreover, taken together, these correlations reinforce the use of *D*_*KL*_(*q*_*i,c*,0_ ∥ *q*_*i,t*_) as a variational proxy for the condition-specific information.

### D. D_KL_ identifies genetic drivers of selection

To characterize any interaction effects systematically associated with selection dynamics across the cancer cell libraries, we used penalized regression to identify the genes that most strongly predicted the *D*_*KL*_ and relative divergence (Fig. 3A and S9A). For all penalties, the cross-validation errors (*CV*_*error*_) were generally lower in the Broad library (Fig. 3B and S9B), with relative divergence being the most predictable (*CV*_*error*_ *<* 0.25 on standardized outcomes). Batch correction with RCPCz eliminated any fitted associations and reduced the number of selected model covariates (Fig. 3B). Interestingly, comparing the fitted associations (model coefficients) with each gene’s average fitness effect (*δ*) revealed how cancer-specific interactions (Δ*δ*) modulate fitness to generate the observed dynamics (Fig. 3A). For overall selection, we observed divergent correlations for genes with positive and negative average fitness, indicating a widening of the distributions for cell environments producing stronger *D*_*KL*_ (Fig. 3B, *upper right panels*). On the other hand, the relative divergence in the Broad data showed a directional modulation, as model coefficients were positively correlated with the average fitness overall (Fig. 3B, *bottom right* panels), further supporting the idea that relative divergence serves as a measure of selective pressure.

**Fig. 3.**
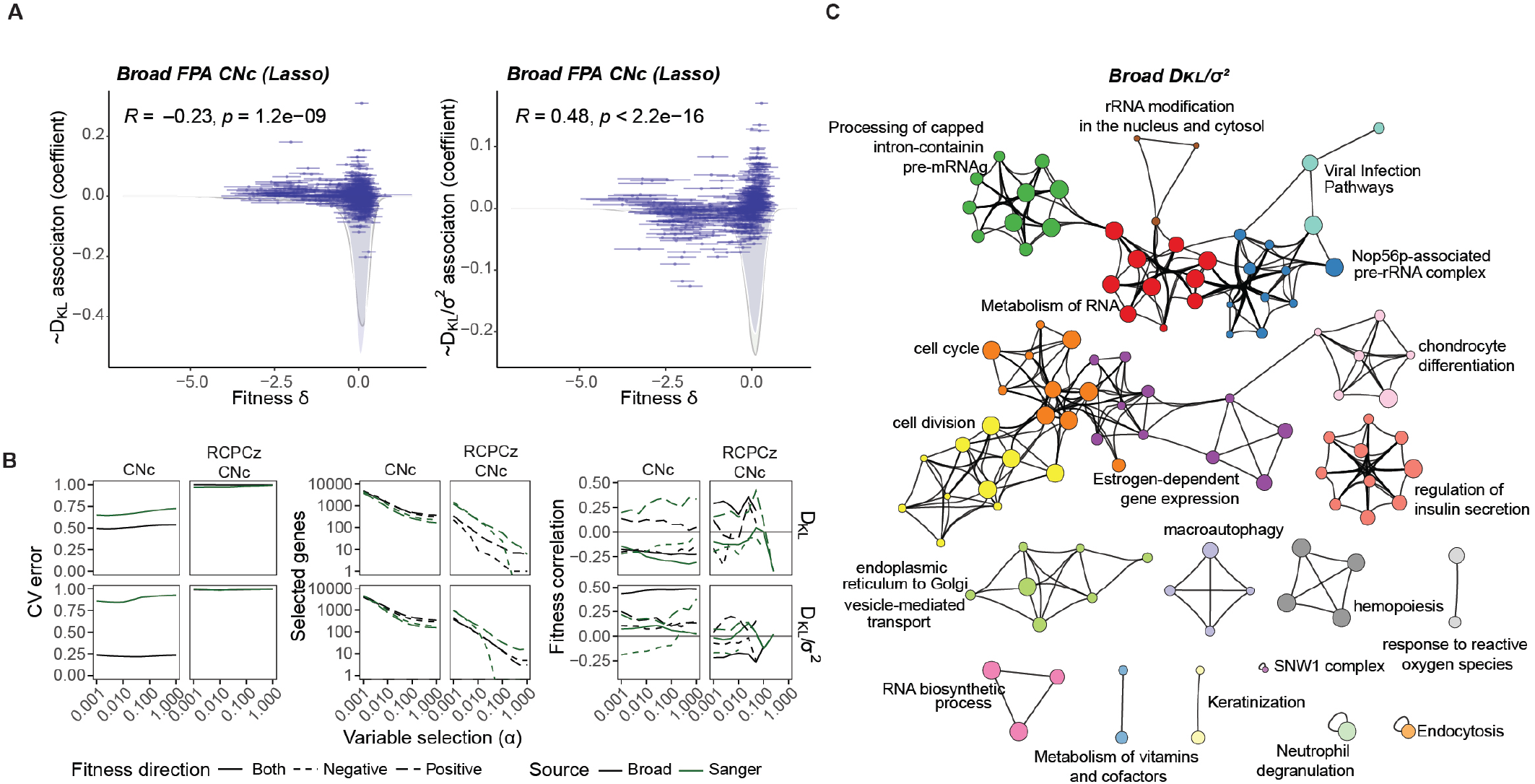
Identification of systematic interaction effects driving selection. **A** Lasso-estimated coefficients for interaction effects in relation to ensemble-averaged fitness perturbations per gene; *R* indicates the Pearson correlation. The fitness distributions show the selected genes (blue shade) compared with the complete genome (gray line). **B** CV error, number of selected genes, and fitness correlations with regression coefficients for *D*_*KL*_ and relative divergence under different variable selection strengths (penalty mixture parameter *α*). **C** Enrichment map for genes whose interaction effects predict relative divergence, where the gene set was selected using Lasso.

We then used Metascape [45] to profile the identified gene sets (Fig. 3C and S9C-E). Two major clusters were associated with the relative divergence in the Broad data—one involving cell cycle regulation, gene expression, and differentiation, and another relating to RNA metabolism and viral infection pathways. These findings suggest that cancer-specific variation in these processes is a major driver of selective pressure in untreated CRISPR screens and a source of batch effects in the pan-cancer libraries.

### E. Generalized fitness perturbation analysis and information theory of selection

The preceding methods analyze each CRISPR screen individually and characterize the functional information gain variationally through *D*_*KL*_(*q*_*i*,0_ ∥ *q*_*i,t*_) and the cellline-specific interaction effects Δ*δ*. To generalize this approach to study information linked to the conditional selection of specific gene mutants, we can reformulate the models to fit an arbitrary environment with fluctuations in conditional dependencies perturbed from a ground state. The change in fitness can be expressed with a finite set (*r* ∈ *R*) of linearly independent perturbation functions such that the growth rate of mutant *i* is 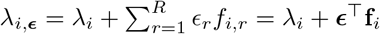, producing the replicator equation 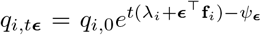. Here, *λ*_*i*_ is the baseline growth rate, while environmental fluctuations in ***ϵ*** define a distinct fitness perturbation profile determined by the gene-specific response parameters **f**_*i*_.

Under this notation, we treat the environmental state as a random variable distributed under *p*(***ϵ***). We can then view fitness optimization through natural selection as a process that reduces uncertainty about environmental fluctuations by strengthening gene-environment couplings (**f**) that increase the probability of a mutant being observed. The functional information encoded in the genome can be quantified using the *mutual information* (see Methods section L), which can be expressed as a change in the conditional distribution of mutants *q*_*i,t****ϵ***_ from an unperturbed distribution *q*_*i,t*_:

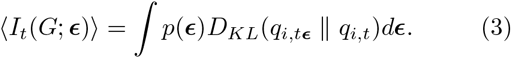

Here, *D*_*KL*_(*q*_*i,t****ϵ***_ ∥ *q*_*i,t*_) = *t****ϵ***^⊤^⟨**f** ⟩_*t****ϵ***_ − Δ*ψ*_***ϵ***_, where 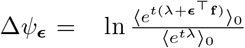 is the joint cumulant generating function.

To quantify the conditional information gain associated with selection, *D*_*KL*_ can be approximated by Taylor expanding ⟨**f**⟩ _*t****ϵ***_ and Δ*ψ*_***ϵ***_ around ***ϵ*** = 0 (or *t* = 0; see Methods section L). Under a second-order approximation

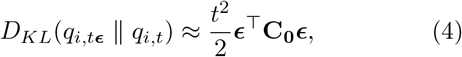

where **C**_**0**_ is the response covariance matrix ⟨ (**f** −⟨**f** ⟩_0_)(**f** − **f**⟩ _0_)^⊤^ ⟩ _*t*_ [46].

Within a linear range, the information gain in response to environmental change can be expressed as ∇ _***ϵ***_*D*_*KL*_(*q*_*i,t*,***ϵ***_ ∥ *q*_*i,t*_) ≈ *t*(⟨**f**⟩ _*t****ϵ***_ −⟨**f**⟩ _0_) ≈ *t*^2^**C**_**0**_***ϵ***. This representation can be viewed as a conditional form of the simple Price equation [4], Δ_*t****ϵ***_ ⟨**f**⟩ = *Cov*[**f**, *tδ*_*t****ϵ***_], which describes the evolution of multivariate traits characterized here by **f**. This also permits expressing the information gain as an expectation, using *t****ϵ*** ≈ **C**_**t**_^−1^(⟨**f** ⟩_*t****ϵ***_ − ⟨**f** ⟩_0_):

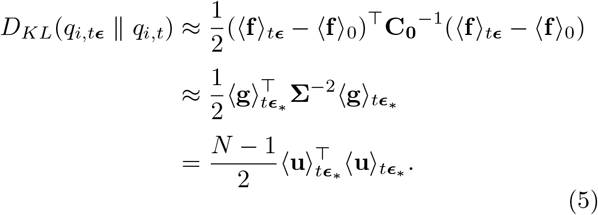

Here, **Σ**^2^ denotes the diagonalized eigenvalues 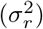 of **C**_**0**_. Meanwhile, 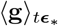 and 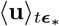 represent the expected principal component scores and normalized projection coordinates (or unscaled component scores), respectively, for the decomposed perturbation functions after selection in environment ***ϵ*** (and its eigentransform ***ϵ***_∗_) (see Methods section L). This approximation offers a simple geometric interpretation of the information gain as the squared distance of normalized projection coordinates in the space of decomposed fitness interactions (Fig. 4A). By marginalizing over *p*(***ϵ***), we obtain mutual information between the environment and the genome after selection: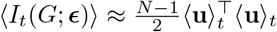.

**Fig. 4.**
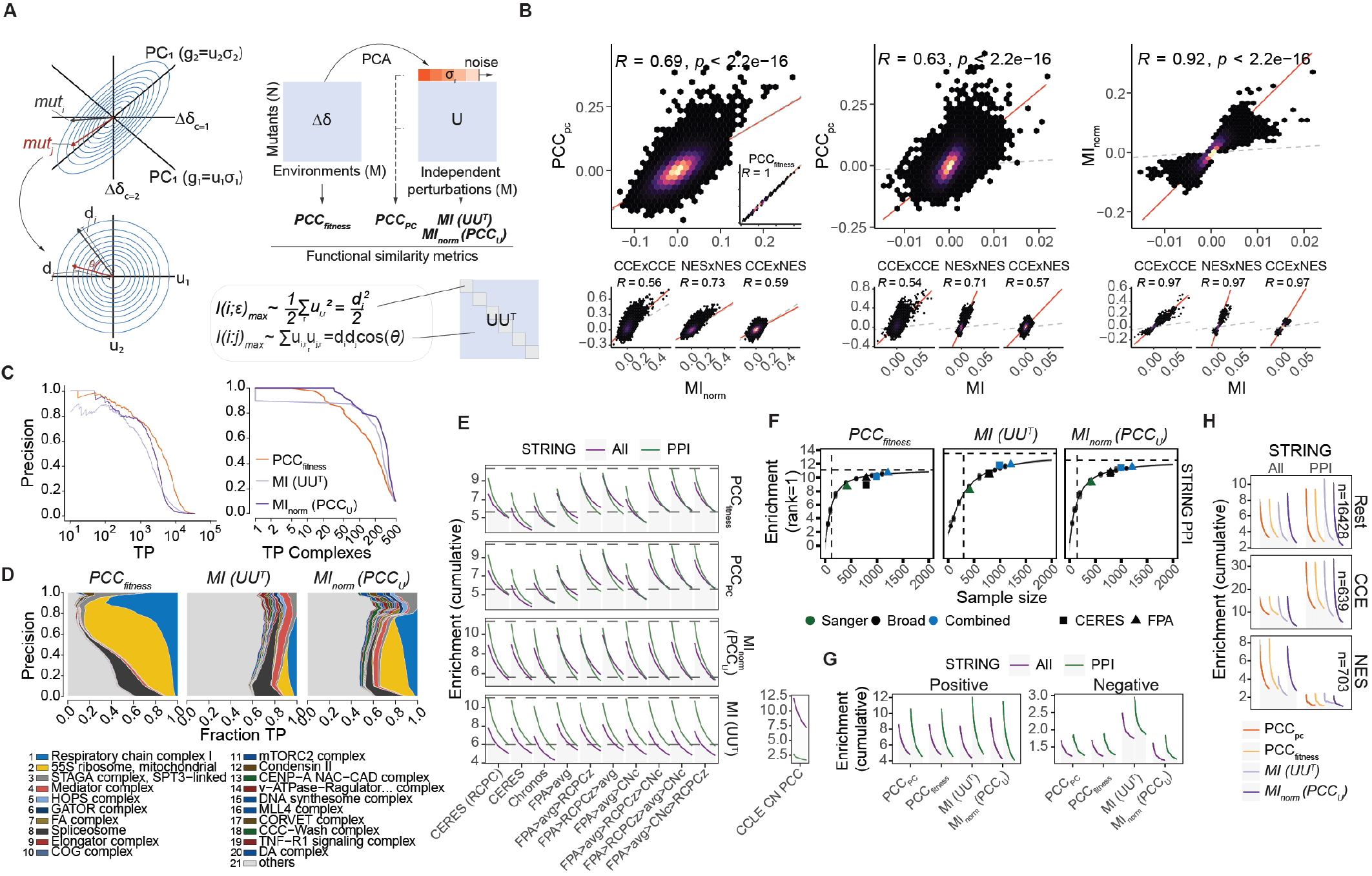
Retrieving gene functional information from decomposed fitness perturbations. **A** Information geometry of gene function (*left*) and overview of functional similarity metrics (*right*). **B** Comparison of similarity metrics. The *top panels* compare random gene pairs (*n* = 100,000), while the *bottom panels* compare gene pairs across and within the CCE and NES gene sets. The inset in the *left panel* shows the correlation between PCC_*PC*_ and PCC_*fitness*_; *R* indicates the Pearson correlation. **C** Gene-level (*left*) and module-level (*right*) precision–recall curves for retrieving CORUM complexes using FLEX. **D** Contribution diversity of CORUM complexes at different precision thresholds. A selected set of complexes is color-coded. **E** Genome-wide enrichment of STRING interactions up to the top 20 rank per functional similarity metric, shown for different fitness computation procedures applied to the Broad Institute dataset. The *lower right* panel shows the enrichment over correlations in CNs. **F** Genome-wide enrichment of STRING PPIs per similarity metric as a function of sample size. The curve represents a Hill equation fitted to enrichment scores from random subsamples (small dots) of the combined FPA datasets. Horizontal and vertical dashed lines indicate the maximum enrichment and EC_50_, respectively. **G** Directional genome-wide enrichment of STRING interactions up to the top 20 rank for each similarity metric, computed from the combined dataset. **H** Genome-wide enrichment of STRING interactions up to the top 50 rank for CCE genes, NES genes, and remaining genome, with similarity metrics computed from the combined dataset.

To analyze the information gain for a specific target state, we can use the *pointwise mutual information* associated with the conditional selection of a mutant. It can be expressed as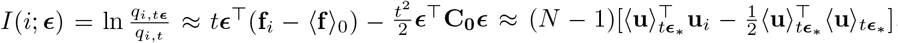. By performing a counterfactual analysis of the perturbations that maximize the selection of mutant *i*, we have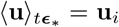, which implies that 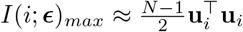. Thus, the diagonal elements of the projection matrix **UU**^⊤^, also known as leverage values, capture the maximal environmental information encoded in each gene (Fig. 4A).

This same logic extends to functional similarities between genes by analyzing the conditions that maximize the co-varying selection of functionally related mutants. We express this through the *pointwise mutual information* shared between mutants: 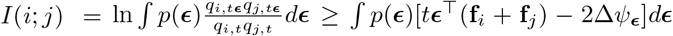 (see Methods section M). Under the second-order approximation and given the perturbations that maximize the selection of a mutant (so that 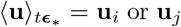), we obtain 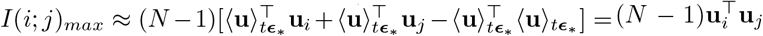. Thus, the off-diagonal elements of the projection matrix, also known as cross-leverage values or influence terms, characterize the maximum functional information shared between any pair of genes (Fig. 4A).

### F. Information-theoretic analysis of CRISPR screen co-selection

The projection matrix **UU**^⊤^ can be generated from a principal component analysis (PCA) of fitness perturbations by performing the SVD: **Δ** = **UD**_∗_**W**^⊤^. Here, **Δ** is the fitness interaction matrix, whose elements correspond to ***ϵ***_*c*_^⊤^ (**f**_*i*_ − ⟨**f**⟩ _0_) = Δ*δ*_*i,c*_ (see Methods section N). Because the rows in **U** are already zero-centered, the Pearson correlations of PCA-based, whitening-transformed fitness scores (*PCC*_*U*_)—as previously described [23]—approximate a normalized mutual information under maximum co-varying selection. This is equivalent to the cosine similarity 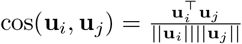 (Fig. 4A).

We next examined these information-theoretic similarity metrics—namely *MI* (corresponding to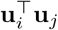) and *MI*_*norm*_*/PCC*_*U*_ (corresponding to cos(**u**_*i*_, **u**_*j*_))—for enrichment of functional gene relations (Fig. 4 and S10). For comparison, we also computed the correlation of fitness perturbations (*PCC*_*fitness*_) and principal components (*PCC*_*PC*_)—with the latter differing from *MI*_*norm*_ by retaining the fitness perturbation magnitudes *σ*_**r**_ (Fig. 4A). This difference could be observed by how similarities for CCE genes (but not NES genes) were relatively stronger under *PCC*_*PC*_ or *PCC*_*fitness*_ compared to *MI*_*norm*_, and less correlated with both *MI* metrics (Fig. 4B). This observation provides an alternative explanation for why fitness correlations are more sensitive in retrieving individual protein pairs from essential CORUM complexes (Fig. 4C, *left*), as noted previously by others [21]. In contrast, complex-level precision-recall was markedly higher for *MI* and *MI*_*norm*_ (Fig. 4C, *right*), showing greater and more uniform complex recall diversity at each precision threshold (Fig. 4D). However, because complexome associations capture only a narrow facet of functional gene relations, pooling the Sanger and Broad datasets yielded only a modest increase in performance (Fig. S10A-C).

As a more appropriate benchmark for genome-wide retrieval of functional relations, we quantified the enrichment of STRING interactions by the gene-wise rank of the similarity metrics [20]. Here, compared to fitness correlations *MI* specifically increased the sensitivity to detect experimentally verified protein-protein interactions (PPIs) but not all STRING interactions (Fig. 4E and S10D). Meanwhile, *MI*_*norm*_ further improved sensitivity for detecting both PPIs and all STRING interactions (Fig. S10D-F), albeit with a faster decline in enrichment per similarity rank. This trade-off suggests that normalizing the gene-wise information content provides greater uniformity and sparsity among significant interactions across the genome.

As noted for similar procedures [20, 23], both *MI* metrics were less sensitive to batch corrections with RCPCz. Meanwhile, FPA with CNc improved the representation of gene functional information, relative to CERES and Chronos, specifically among fitness correlations (Fig. 4E and S10E-G). For *MI*-based metrics, Chronos performed substantially worse than CERES and FPA, suggesting a permanent loss of functional information. Interestingly, analyzing the contribution of each correction procedure in FPA revealed a CN-associated detection bias for non-PPIs in uncorrected data (Fig. 4E, *lower right panel*).

Computing similarity scores under subsampling of the combined dataset showed that while PPI enrichment from *PCC*_*fitness*_ was near its asymptotic maximum, *MI* had substantially more to gain with increasing sample size (Fig. 4F and S10H). Furthermore, *MI* showed much higher PPI enrichment for negative gene associations—rare in all correlation-based metrics (Fig. 4G)—indicating an asymmetry in information content distribution among antagonistic gene pairs. Lastly, grouping by gene essentiality revealed that both *MI* metrics selectively improved PPI enrichment for non-core essential genes compared to fitness correlations (Fig. 4H).

### G. Functional enrichment across the MI landscape

To characterize the functional structures that differentiate *MI*-based metrics from fitness-based similarities, we systematically performed a series of GO enrichment tests (Fig. 5 and S11). Although it correlated strongly with fitness variance, the maximum information content per gene 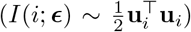 yielded stronger enrichment across all GO libraries when assessed using GSEA (Fig. 5A and S11A-B) [47]. Moreover, comparing the GO terms with selectively higher enrichment for each metric revealed that metabolism and mitochondrial activity were more associated with higher fitness variance, whereas processes related to chromatin remodeling, transcription, and DNA replication were linked to higher information content (Fig. 5A).

**Fig. 5.**
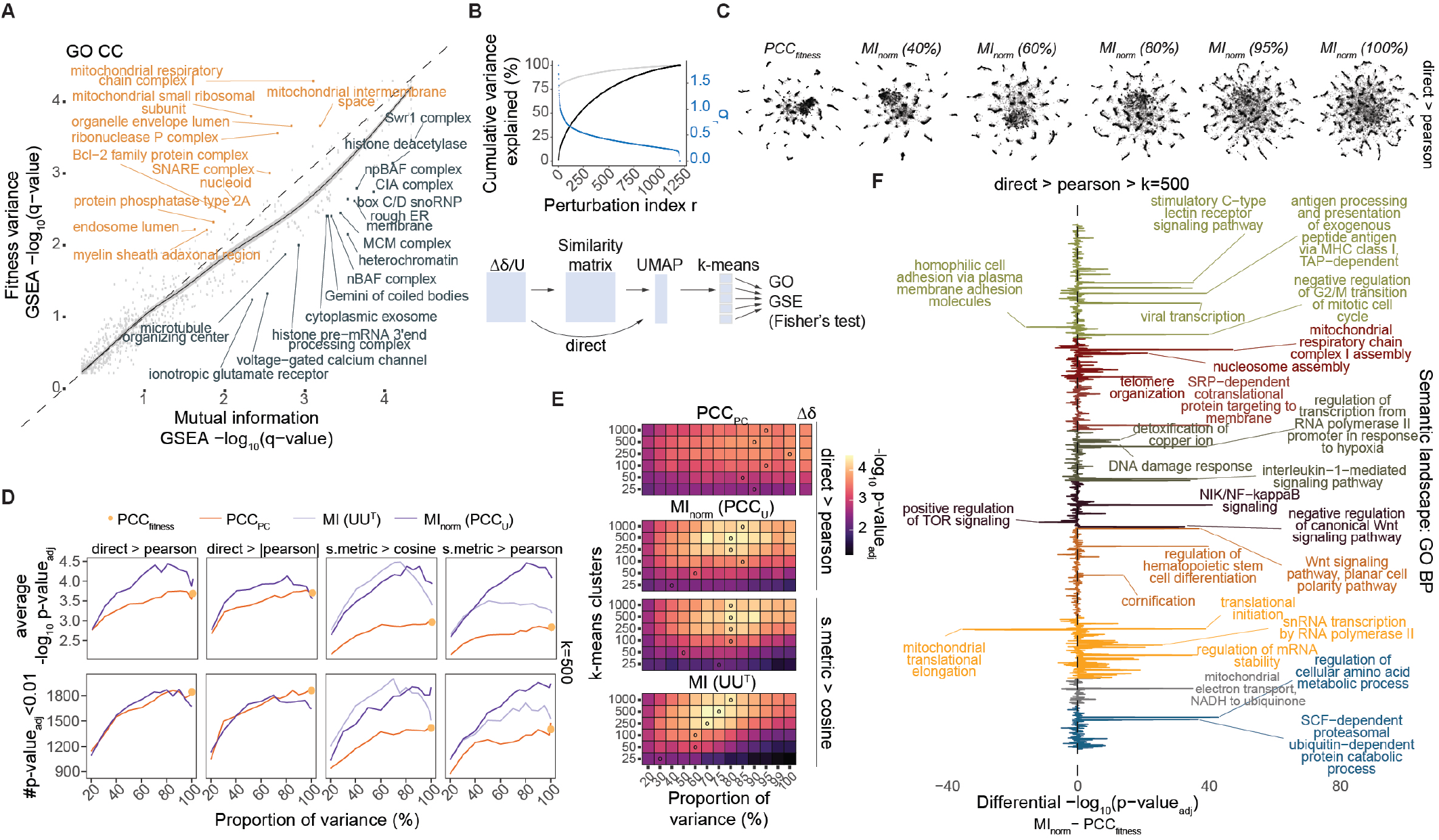
Functional associations and optimal embedding of the MI landscape. **A** GSEA GO-CC enrichment for *I* (*i*; ***ϵ*)** and 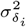. A LOESS regression line with a 95% CI is shown. **B** (*Top panel*) Cumulative variance explained, and (*bottom panel*) an overview of the UMAP embedding of functional similarity metrics. Black and gray lines indicate the cumulative variance explained (%) for decomposed Δ*δ* and *δ*, respectively, while the blue line shows the variance per principal component for Δ*δ*. **C** UMAP embedding of genome-wide functional landscapes, showing *PCC*_*fitness*_ and *MI*_*norm*_ generated by embedding **Δ***δ* and **u** using Pearson correlation as the distance metric. Different perturbation ranks for *MI*_*norm*_ correspond to the cumulative proportions of variance indicated in B. **D** GO term enrichment summaries (average − log_10_ (*p*-value_adj_), *top panels*; number of significant enrichments, *bottom panels*) for different UMAP embedding configurations; *k*-means clusters *k* = 500. **E** Average GO term enrichment statistics for varying *k*-means clusters and different perturbation ranks, each corresponding to cumulative proportions of variance indicated in B. **F** Differential enrichment per GO-BP term comparing *PCC*_*fitness*_ and *MI*_*norm*_, using a perturbation rank corresponding to 80% of the variance.

To investigate the global architecture of this mutual information landscape, we used Uniform Manifold Approximation and Projection (UMAP) to embed the similarity data in a two-dimensional graph [48]—either directly from fitness perturbations (Δ*δ*) and the projection coordinates (*U*), or by applying a second distance transform to the similarity matrices (Fig. 5B). We then stratified the embedded graphs using k-means clustering with various thresholds and assessed them for enriched GO-BP terms via Fisher’s exact test.

Observing a trade-off between precision and PPI enrichment depth for *MI* and *MI*_*norm*_ at different perturbation ranks (Fig. S10F), we hypothesized that adjusting the number of included perturbations could tune the resolution of the embedded landscape (Fig. 5B). For reference, both *PCC*_*fitness*_ and *PCC*_*PC*_ displayed more granular embedding (Fig. 5C and S11C-D). Notably, reducing the perturbation rank for *PCC*_*PC*_ only diminished GO term enrichment (Fig. 5D). In contrast, increasing the perturbation rank made *MI*-based embeddings more fine-grained, with the landscape granularity shifting alongside the k-means cluster threshold to achieve optimal GO enrichment (Fig. 5D). Moreover, detection of functional information in smaller clusters peaked at a perturbation rank corresponding to roughly 80% of the variance explained (Fig. 5E, S11E), greatly improving GO term enrichment across the entire GO-BP landscape (Fig. 5F, S11F).

By applying a quantile-based cutoff (*p ≤* 0.1%; Fig. S12A) to the similarity scores, we generated functional networks and overlaid them onto the UMAP embedding to analyze the cell’s functional network structure and connectivity (Fig. 6). We then used Spatial Analysis of Functional Enrichment (SAFE) to label unique network domains across the UMAP [49]. This procedure revealed a remarkable resolution of *MI*_*norm*_ embedded networks when using a secondary Pearson transform of the absolute *MI*_*norm*_-values (Fig. 6A).

**Fig. 6.**
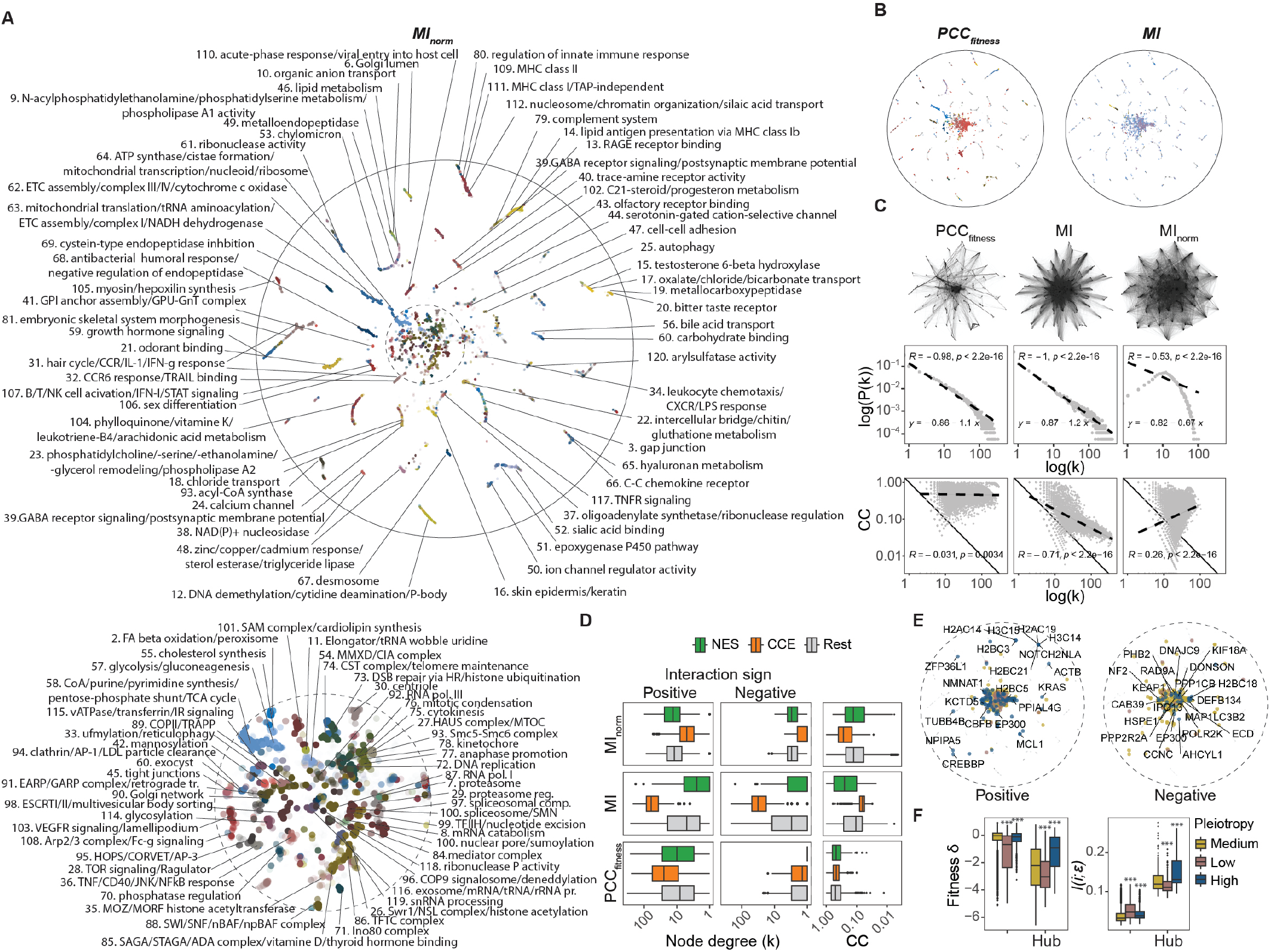
Genome-wide functional network structure. **A** *MI*_*norm*_ SAFE network domain annotation. The landscape represents a UMAP of absolute *MI*_*norm*_ values, embedded using Pearson correlation as the distance metric. The *top panel* shows the genome-wide architecture, and the *bottom left panel* shows a magnified view of the central core. **B** *PCC*_*fitness*_ and *MI* SAFE network domain annotations, visualized over the *MI*_*norm*_ embedding from panel A. **C** *PCC*_*fitness*_, *MI*, and *MI*_*norm*_ networks, each visualized over the *MI*_*norm*_ embedding (*top panels*). The node degree distributions are shown as log–log plots to assess power-law scaling (*middle panels*), and clustering coefficients (CC) are plotted against node degrees to evaluate network structure (*bottom panels*). *R* indicates the Pearson correlation; linear fits (dashed lines) are provided. **D** Node degrees stratified by edge sign and clustering coefficients (CC) for CCE genes, NES genes, and the remainder of the genome. **E** Distribution of *MI* gene hubs, stratified by edge sign. **F** Average fitness *δ* and *I* (*i*; ***ϵ*)** for *MI* gene hubs, with color coding in panels E and F indicating pleiotropy.

In contrast to previous reports [20], these maps exhibited a progressive distribution of tissue-specialized functions spanning immunity, migration, metabolism, and neuronal signaling across cortical clusters, as well as a highly resolved core of cellular functions in the graph’s center (Fig. 6A). Closer inspection of the core showed an organized compartmentalization of functionally related substructures (Fig. 6A, *lower panel*), annotated with an accuracy previously observed only in simpler genomes of lower eukaryotes [6].

In comparison, *PCC*_*fitness*_ and *MI* networks provided more limited annotations (Fig. 6B and S12B-C). *PCC*_*fitness*_ primarily annotated core cell processes as a single unresolved domain alongside an over-connected mitochondrial cluster (Fig. 6B-C). On the other hand, *MI* produced a highly hierarchical network structure, featuring genome-wide connections radiating from the cell core (Fig. 6C). Analysis of network connectivity revealed that both *PCC*_*fitness*_ and *MI* networks followed a power-law distribution (Fig. 6C), though they were not fully scale-free (scale parameter *<* 2) [50, 51]. Notably, *MI* showed a modular structure containing an overconnected essential gene subgraph (Fig. 6C-D and S13A). In comparison, the *MI*_*norm*_-based network displayed a degree distribution and clustering coefficients consistent with a strong uniform community structure (Fig. 6C), wherein essential gene subgraphs were less dense but highly interconnected (Fig. 6D and S13A). This inversion of *MI* and *MI*_*norm*_-network structures reflects the normalization of highly information-rich essential cell processes.

An analysis of *MI* network hubs revealed a disproportionate enrichment in the center (Fig. 6E), featuring genes with more negative average fitness perturbations (i.e., more essential) and higher mutual information (Fig. 6F). Moreover, classifying network hubs by metric directionality and pleiotropy (wiring distance variability) showed that regulatory hub genes with high pleiotropy for negative interactions were exclusively located in the functional core. Notably, although pleiotropic hub genes had higher mutual information, they were less essential for growth than low-pleiotropy hubs (Fig. 6F), suggesting that multifaceted regulators exhibit more complex fitness perturbation profiles (Fig. S13B). This finding also indicates that high information content manifests itself not only in high connectivity but also in wiring variation across diverse functional domains. Charting these differences may be crucial for understanding the robustness, vulnerability, and unpredictability of biological systems that respond to perturbations.

### H. Information gain in cell-type-specific subspaces

A key challenge in studying the functional architecture of the human genome lies in the complexity of contextdependent regulation and the rewiring of functional relationships in cells of different tissue origins [25, 52]. This rewiring shapes the immediate environment that modifies genetic dependencies, which should be reflected in the space of possible fluctuations of ***ϵ***. To explore this further, we leveraged the complementarity between the decomposed perturbations in **U** and **W** by removing a subset of cell types *S* from **W** and rotating the components of the projection matrix using 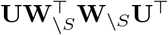. Thus, cell-type-specific information under maximum selection was defined as the difference between native and rotated *MI* scores (Δ*MI*; see methods).

Applying a quantile-based cutoff (*p* ≤ 0.1%) to Δ*MI*_*norm*_ revealed cell-type-specific networks, uncovering functional clusters absent from the native genomewide network (Fig. 7A). This was evident when comparing *B* and *T cells*, which enriched for B and T cell receptors, respectively, along with immune cell-specific functions, signaling modules, and differentiation factors (Fig. 7A). *Neurons*, representing a highly specialized cell phenotype, presented a more intricate network with clusters related to nervous system development, synaptic vesicle proteins, and action potential (Fig. S14A). Notably, brain developmental factors such as ASCL1, SOX11, NEUROD1, PHOX2A, and PHOX2B emerged as major hub genes within the network.

**Fig. 7.**
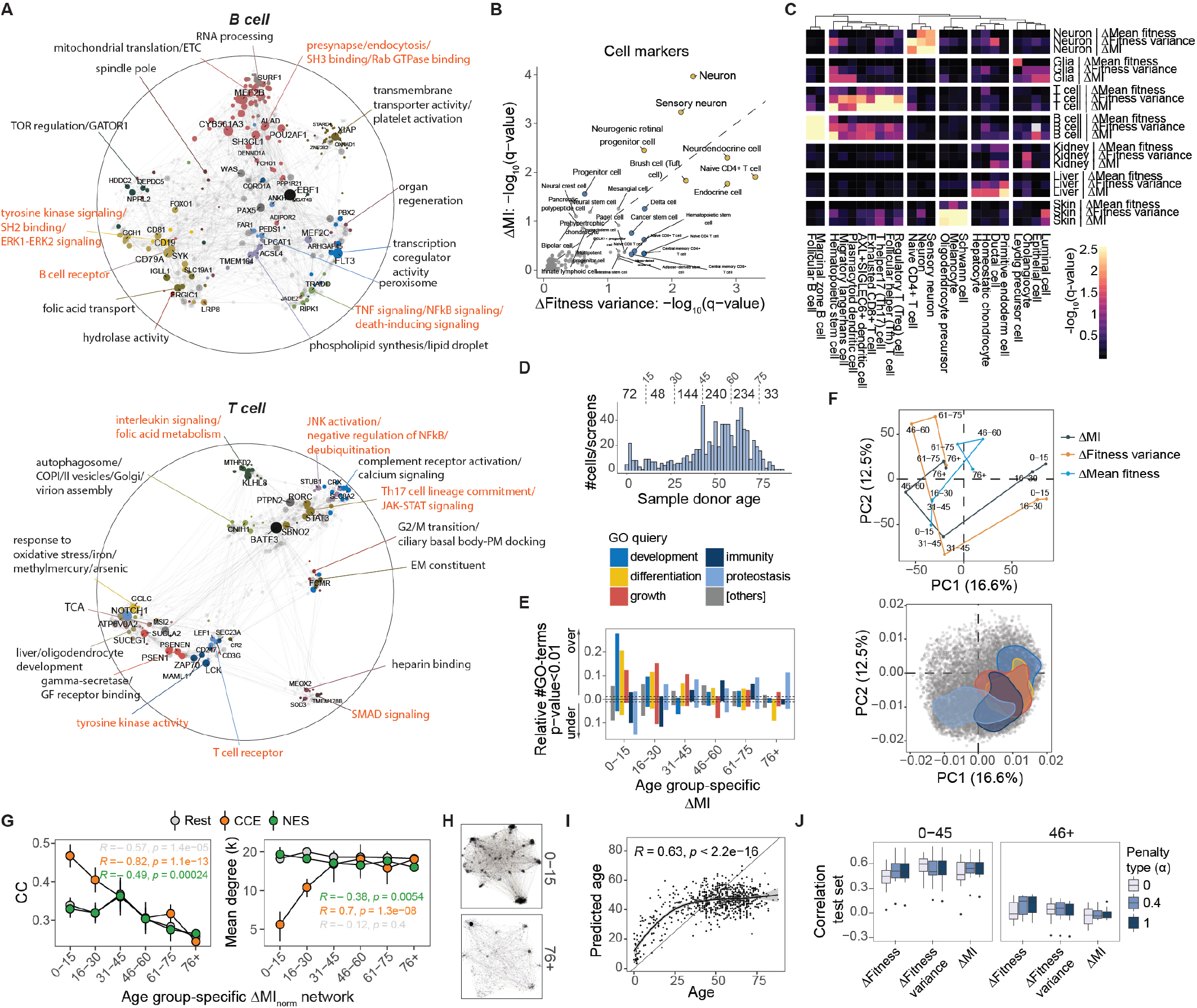
Cell-type-specific information captures functional signatures of tissue differentiation, development, and aging. **A** Δ*MI*_*norm*_ SAFE network domain annotation. The landscapes represent UMAPs of absolute Δ*MI*_*norm*_ values, embedded using Pearson correlation as the distance metric. Labels in red indicate cell-type-specific domains. **B** GSEA cell-type marker enrichment for Δ*I* (*i*; ***ϵ*)** and 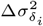 in neurons. Yellow and blue highlight commonly or uniquely significant terms, respectively. **C** Cell-type marker enrichment summary, showing the top three terms for each respective cell type and metric. The heatmap scale is capped at − 2 log_10_(0.05). Hierarchical clustering was performed using Euclidean distances and complete linkage. **D** Distribution of reported sample-donor ages for the screened cell lines. **E** Sensitivity of significant GSEA enrichment per indicated GO query for Δ*I* (*i*; ***ϵ*)** across different age groups. Directions indicate over- or under-representation. **F** PCA of normalized enrichment scores (NES) from **E**, showing principal component scores per age group (*top*) and PCA loadings where the distribution contours of the indicated GO queries are outlined (*bottom*). **G** Clustering coefficient (CC) and mean node degree **G**.for CCE genes, NES genes, and the remaining genome in age-group–specific Δ*MI*_*norm*_ networks. Points and bars represent the means and standard deviations from bootstrapped age-group subsets of size *b* = 33, ensuring equal sample sensitivity across age groups; *R* indicates the Spearman correlation. **H** Age-group–specific Δ*MI*_*norm*_ networks, embedded as in panel *A*. **I** Lasso-penalized linear regression of age from cell-line–specific Δ*I* (*i*; ***ϵ*)** values; *R* indicates the Pearson correlation comparing predicted versus true donor age, and the curve represents a LOESS fit, illustrating the smoothed average prediction by age. **J** Correlation of 10-fold test predictions of age (log) per indicated age group for penalized linear models trained on different cell-line–specific metrics.

GSEA enrichment of cell markers [53] across different gene-wise differential scores revealed that Δ*MI* matched cell type and cell markers with higher sensitivity compared to differences in fitness variance 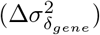 or mean fitness 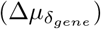 (Fig. 7B-C and S14B). This pattern extended to cell types with less distinct characteristics, enriching for markers of functionally similar cells. Moreover, GO-BP enrichment of *neurons* highlighted that cell-type-specific functions ranked among the top terms for Δ*MI* but not for differential fitness associations (Fig. S14B).

Finally, to assess whether Δ*MI* could capture functional differences across human development and aging, we grouped cells by donor age and performed GSEA for these defined age groups (Fig. 7D). Remarkably, the detection sensitivity for GO terms related to *development, differentiation*, and *growth* changed gradually across age groups, peaking sequentially during the early stages of life (Fig. 7E). In contrast, GO terms related to *immunity* and *proteostasis* were under-represented in early life but gradually increased in detection sensitivity in older age groups. These transitions were more pronounced for Δ*MI* (Fig. S14C) and aligned with a temporal grouping of age clusters when applying PCA to enrichment scores across the entire GO library (Fig. 7F).

As overall enrichment declined with age, we speculated whether age-specific Δ*MI* data could reflect traits of biological aging. Analysis of age-group–specific network structures revealed a significant decline in Δ*MI*_*norm*_ network clustering with age, along with relative changes in connectivity among CCE and NES genes (Fig. 7G-H and S14D). These changes also influenced screen selection dynamics, with increased noise present in cells from older individuals (Fig. S14E-F). Finally, while differences in chronological age could be predicted during early life, this was not possible for individuals over 45 years (Fig. 7I-J and S14G-I). This outcome indicates that time-dependent functional rewiring characterizes development but not aging in human cells. Altogether, these results highlight the advantages of the presented information-theoretic framework in analyzing gene function, offering unique insights into development and cell-type-specific behavior.

## III. DISCUSSION

Since the discovery of the genetic code, there has been a significant interest in formulating evolution in the language of information theory. The goal is to quantify the information that organisms gain from the environment through natural selection acting on biological functions. In this context, the concepts of information and function are intimately linked. However, to date, evolutionary change has primarily been studied through the concept of fitness effects, where the evolvability of a trait is quantified using its fitness covariance, as formalized by the Price equation [4]. Information-theoretic extensions have been developed using the *D*_*KL*_ to compare populations before and after selection [30]. However, these metrics represent only a single random variable (*n* = 1), and without strong contextual data, they offer limited insight into the utility of the selected trait. These challenges also encompass well-known pitfalls of applying causal interpretations to the Price equation without additional modeling assumptions [5], as well as complexities arising from pleiotropy and context-dependent traits [2].

Here, we used pooled CRISPR screens as an experimental study system to develop a coherent framework to quantify the functional information content associated with variational selection of well-defined genotypes (i.e., gene deletions). Surprisingly, the application of an exponential growth model with minimal assumptions challenged many standard methods for CRISPR screen analysis across both conventional and novel benchmarks. Consistent with model predictions, our analyses further reveal that the selection dynamics is largely captured by the first three cumulants of the fitness distribution and, in the absence of applied selection pressure, display characteristic asymptotic behavior tending towards a parabolic relationship between skewness and kurtosis [39].

Based on these results, we proposed using the *D*_*KL*_ and its variance-normalized form (relative divergence) to quantitatively assess information gain and selective pressure. For example, the distinct selection dynamics induced by low-dose drug treatment (Fig. 1H) were identified through the relative divergence but not by fitness variance, skewness, or kurtosis individually (Fig. S2B,E). In CRISPR screen ensembles, this metric identified novel gene-environment interactions driving selection and masking the detection of other fitness associations. These findings differ from the sample time-specific batch effect observed when comparing screens from the Broad and Sanger Institutes [21]. Although RCPCz mitigates this effect by altering library-wide fitness correlations (Fig. S4E–F), it highlights the importance of considering potential confounding effects from cell-specific properties interacting with experimental parameters [54, 55]. Furthermore, when analyzing bidirectional functional associations through Δ*δ, D*_*KL*_ proved to be an informative metric even after batch correction (Fig. S6 and S8). Our framework establishes a direct link between condition-specific information *I*_*t*_(*G, C* = *c*) and Δ_*c*_*D*_*KL*_(*q*_*i*,0_ ∥ *q*_*i,c,t*_), enabling the variational use of the *D*_*KL*_ in cases where baseline conditions are difficult to define.

An additional advantage of the presented modeling framework is its capacity to directly link information-theoretic measures to explicit interaction parameters of the fitness distribution [46]. In this way, function—quantified from selection paths in which mutual information between genetic and environmental variables is maximized—can be analyzed within a straightforward geometric framework utilizing normalized projection coordinates. These findings not only provide practical heuristics that bridge theoretical concepts with experimental data, but also align with formal derivations of *D*_*KL*_ as a generalized squared distance for classes of exponential family distributions [56]. Interestingly, the presented method also links differential information gain with a multivariate expression of the simple Price equation. However, beyond serving as a statistical identity for evolutionary change [4, 5], this framework requires counterfactual reasoning about environmental conditions that would maximize gene function selection. This introduces a more generative perspective to gene functional profiling [25], where context-dependent traits are selected within geometric subspaces, revealing the dynamic rewiring of functions across the genome.

Although the effects of covariance whitening have previously been studied [23], our work shows that this metric is a normalized form of the pointwise mutual information (*MI*_*norm*_) and should not be interpreted as a coessentiality score [20]. In fact, *MI*_*norm*_ demonstrated reduced sensitivity to detect functional relationships involving essential genes (Fig. 4B,H). However, likely due to its greater uniformity, *MI*_*norm*_ proved superior for embedding complex functional manifolds on a genome-wide scale compared to *MI* and *PCC*_*fitness*_ (Fig. 5-6). Conversely, *MI* captured the hierarchical structure of the cell and facilitated deeper retrieval of functional gene pairs linked to core regulatory network hubs. *MI* also enabled a broader assessment of the genomic distribution of functional information overall (Fig. 5A and 6E-F) and differential information across cell environments (Fig. 7).

Finally, an analysis of sample size dependencies revealed that *MI* exhibited greater gains with increasing numbers of screened cell lines (Fig. 4F). This finding highlights the potential to enhance the latent data structure and further deepen the functional annotation of the human genome. Incorporating greater cell lineage variability and integrating transcriptomic data to explore how differential gene expression shapes local dependencies could provide new opportunities to reverse engineer the functional architectures underlying human cell types and their behavior.

## AUTHOR CONTRIBUTIONS

Conceptualization: A.N.A and J.M.E. Writing - first draft: A.N.A. Writing - editing: A.N.A., N.C., and J.M.E. Figures: A.N.A. and N.C. Data analysis: A.N.A and N.C. Modeling and statistics: A.N.A. Experiment: L.P. Software: A.N.A. and S.N. Supervision: M.Z. and J.M.E. Funding: J.M.E. All authors reviewed the manuscript.

## ACKNOWLEDGMENTS

This work was supported by grants from the Norwegian Health Authority South-East (project numbers 2018012 and 2025077), The Norwegian Cancer Society (project number 208012), and the Norwegian Research Council (project number 314811). This work was partly supported by the Research Council of Norway through its Centres of Excellence funding scheme (project number 262652).

## Notes

### Competing Interest Statement

The authors have declared no competing interest.

