## Supplemental methods and figures. for "Perturbational fitness analysis of CRISPR screens uncovers information-theoretic relation between gene function and selection"

### IV. SUPPLEMENTARY METHODS

#### A. Computing fitness perturbations from sgRNA sequence reads

The replicator equation measures the change in frequency $q_i$ of a mutant $i$ in a mixed population of mutants where $q_i = \frac{n_i}{\sum_i n_i}$ and $n_i$ is the cell number. If each mutant is growing exponentially with a constant growth rate, the cell number at time $t$ is $n_{i,t} = n_{i,0}e^{t\lambda_i}$, and $q_{i,t} = \frac{n_{i,0}e^{t\lambda_i}}{\sum_i n_{i,0}e^{t\lambda_i}} = q_{i,0}\frac{e^{t\lambda_i}}{\langle e^{t\lambda_i}\rangle_0} = q_{i,0}e^{t\lambda_i-\psi}$. $\langle\cdot\rangle_0$ indicates the expectation at $t = 0$. The cumulant generating function $\psi = \ln\langle e^{t\lambda_i}\rangle_0 = \sum_{k=1}^{\infty}\frac{t^k}{k!}\frac{d^k\psi}{dt^k}|_{t=0} = \sum_{k=1}^{\infty}\frac{t^k}{k!}\kappa_{\lambda,k}$. The first four cumulants are $\kappa_{\lambda,1} = \langle\lambda\rangle_0 = \mu_\lambda$, $\kappa_{\lambda,2} = \langle(\lambda-\mu_\lambda)^2\rangle_0 = \sigma_\lambda^2$, $\kappa_{\lambda,3} = \langle(\lambda-\mu_\lambda)^3\rangle_0$, and $\kappa_{\lambda,4} = \langle(\lambda-\mu_\lambda)^4\rangle_0$. Combining $\ln\frac{q_{i,t}}{q_{i,0}} = t\lambda_i - \psi$ with $D_{KL}(q_{i,0} \parallel q_{i,t}) = \langle\ln\frac{q_{i,t}}{q_{i,0}}\rangle_0 = t\mu_\lambda - \psi$ we get the solution in equation 2.

For sgRNA sequence reads $x_{i,t}$, the mutant frequencies were calculated as $q_{i,t} = \frac{x_{i,t}}{X_t}$, where $X_t = \sum_i x_{i,t}$ is the sequencing depth for a given sample time. To address zero counts at later time points, we applied Laplace smoothing (+1 pseudocount): $\hat{q}_{i,t} = \frac{q_{i,t}X'+1}{\sum_i(q_{i,t}X'+1)}$, where $X'$ denotes the lowest sequencing depth in the screen, or a gradient smoothing $\hat{q}_{i,t} = \frac{q_{i,t}X'+1mr_t}{\sum_i(q_{i,t}X'+1mr_t)}$, where $mr_t = \frac{median(\boldsymbol{q_t})}{median(\boldsymbol{q_0})}$.

For time series, the fitness perturbation $\delta_i = \lambda_i - \mu_\lambda$ was estimated using linear regression for each sgRNA identity $i$, with $\ln\frac{\hat{q}_{i,t}}{\hat{q}_{i,0}} + \langle\ln\frac{\hat{q}_{i,t}}{\hat{q}_{i,0}}\rangle_0 = t\beta_i + e_{i,t}$, where $\delta_i \approx \beta_i$.

Alternatively, fitness perturbations were computed by spectral decomposition. With the matrix $\boldsymbol{R} = [\boldsymbol{r}_{t_1}, \boldsymbol{r}_{t_2}, ..., \boldsymbol{r}_{t_f}]$ containing the elements $r_{i,t} = \frac{\hat{q}_{i,t}}{\hat{q}_{i,0}} \approx e^{t\lambda_i-\psi}$, the growth rates $\boldsymbol{\lambda} = [\lambda_1, \lambda_2, ..., \lambda_N]$ define the initial expansion of the system and scale with the first left-singular vector as $\boldsymbol{\lambda} \sim \frac{1}{\tau}\ln|\boldsymbol{a}_{(1)}|$, from the singular value decomposition $\boldsymbol{R} = \boldsymbol{ADB}^\top$, where $\tau$ is the average sample time. The fitness perturbations can be estimated as $\delta_i \approx \frac{1}{\tau}(\ln|\boldsymbol{a}_{(1)}| - \langle\ln|\boldsymbol{a}_{(1)}|\rangle_0)$. For estimates from a single time points this approach yielded the exact same spectrum as equation (2). However, for all SVD-based FPA results in Fig. S2, $\ln|\boldsymbol{a}_{(1)}|$ was centered around the median instead of the $q_{i,0}$-weighted expectation.

#### B. Evaluation of selection dynamics using the $\mathbf{D_{KL}}$

From $\ln\frac{q_{i,t}}{q_{i,0}} - \langle\ln\frac{q_{i,t}}{q_{i,0}}\rangle_0 = t(\lambda_i - \mu_\lambda)$, the higher-order terms of $\psi$ were estimated as $\frac{t^2}{2}\sigma_\lambda^2 = \frac{1}{2}\sigma_{\ln r_{i,t}}^2$, $\frac{t^3}{6}\kappa_{\lambda,3} = \frac{1}{6}\sigma_{\ln r_{i,t}}^3 S_{\ln r_{i,t}}$ with the standardized skewness $S_{\ln q_t/q_0} = \langle(\frac{\ln r_{i,t}-\mu_{\ln r_{i,t}}}{\sigma_{\ln r_{i,t}}})^3\rangle$, and $\frac{t^4}{24}\kappa_{\lambda,4} = \frac{1}{24}\sigma_{\ln r_{i,t}}^4 K_{lnr_{i,t}}$ with the standardized kurtosis $K_{\ln r_{i,t}} = \langle(\frac{\ln r_{i,t}-\mu_{\ln r_{i,t}}}{\sigma_{\ln r_{i,t}}})^4\rangle$. Evaluation of the CRISPR screen selection dynamics was performed by computing the errors or correlations between $D_{KL}(q_{i,0} \parallel q_{i,t})$ and the step-wise combination of $\frac{1}{2}\sigma_{\ln r_{i,t}}^2$, $\frac{1}{6}\sigma_{\ln r_{i,t}}^3 S_{\ln r_{i,t}}$, and $\frac{1}{24}\sigma_{\ln r_{i,t}}^4 K_{\ln r_{i,t}}$. This was also done between the relative divergence $2D_{KL}(q_{i,0} \parallel q_{i,t})/\sigma_{\ln r_{i,t}}^2$ and the step-wise combination of 1, $\frac{1}{3}\sigma_{\ln r_{i,t}} S_{\ln r_{i,t}}$, and $\frac{1}{12}\sigma_{\ln r_{i,t}}^2 K_{\ln r_{i,t}}$. Estimation of the Taylor coefficients was done by fitting the model $D_{KL}(q_{i,0} \parallel q_{i,t}) \sim \sigma_{\ln r_{i,t}}^2 + \sigma_{\ln r_{i,t}}^3 S_{\ln r_{i,t}} + \sigma_{\ln r_{i,t}}^4 K_{\ln r_{i,t}}$. To find the best model explaining the variation in $D_{KL}$, we included the higher-order cumulants in a step-wise manner and scored the model fits using the Bayesian information criterion (BIC) and $R^2$.

#### C. Extensions on the $\mathbf{D_{KL}}$

$D_{KL}(q_{i,0} \parallel q_{i,t}) = \langle\ln\frac{q_{i,0}}{q_{i,t}}\rangle_0 \geq 0$ will measure the information gain associated with negative selection of mutants, as the center of the distribution $q_t$ moves towards zero. If the distribution $q_0$ is uniform, this gives a balanced measure of the average selection pressure. $D_{KL}(q_{i,t} \parallel q_{i,0}) = \langle\ln\frac{q_{i,t}}{q_{i,0}}\rangle_t \geq 0$ will measure information gain associated with the positive selection of mutants that grow $q_t$ relative to their initial frequency $q_0$. Similarly to $D_{KL}(q_{i,0} \parallel q_{i,t}) = -t\langle\lambda_i\rangle_0 + \ln\langle e^{t\lambda_i}\rangle_0 = \frac{t^2}{2}\sigma_\lambda^2 + \frac{t^3}{6}\kappa_{\lambda,3}...$, we can analyze $D_{KL}(q_{i,t} \parallel q_{i,0}) = t\langle\lambda_i\rangle_t - \ln\langle e^{t\lambda_i}\rangle_0$. By Taylor expanding $\langle\lambda_i\rangle_t$ around zero such that $t\langle\lambda_i\rangle_t = t\mu_\lambda + t^2\sigma_\lambda^2 + \frac{t^3}{2}\kappa_{\lambda,3}...$, we get

$$D_{KL}(q_{i,t} \parallel q_{i,0}) = \frac{t^2}{2}\sigma_\lambda^2 + \frac{t^3}{3}\kappa_{\lambda,3}.... \quad (6)$$

Thus, we can see that $D_{KL}(q_{i,t} \parallel q_{i,0})$ as a function of the fitness distribution cumulants at $t = 0$ is closely related to $D_{KL}(q_{i,0} \parallel q_{i,t})$, but expands with $k$th order coefficients equal to $\frac{1}{(k-2)!k}$ compared to $\frac{1}{k!}$. This gives relatively more weight to the higher-order cumulants (Fig. S4C, *top panels*), which reflects how $D_{KL}(q_{i,t} \parallel q_{i,0})$ will measure the information gain associated with positive selection, as the right tail of the distribution $\lambda_i$ outgrows the remaining population. This divergence is also capped by an upper bound $\ln\frac{1}{q_{j,0}}$ as $q_{j,t} \to 1$ for the winning replicator $j$.

The symmetrized $D_{KL}$, or Jeffreys divergence, which has previously been related to the average change in fitness [4], can now be described more generally as

$$\begin{aligned} J(q_{i,t}, q_{i,0}) &= D_{KL}(q_{i,t} \parallel q_{i,0}) + D_{KL}(q_{i,0} \parallel q_{i,t}) \\ &= t(\langle\lambda_i\rangle_t - \langle\lambda_i\rangle_0) = t\Delta_t\langle\lambda_i\rangle \\ &= t^2\sigma_\lambda^2 + \frac{t^3}{2}\kappa_{\lambda,3}..., \end{aligned} \quad (7)$$

with $k$th order coefficients equal to $\frac{1}{(k-1)!}$. Thus, under the assumption of a fixed environment and exponential growth [57], the rate of change in average fitness

$$\frac{d\langle\lambda_i\rangle}{dt} = \sigma_\lambda^2 + \frac{t}{2}\kappa_{\lambda,3}..., \quad (8)$$

extends Fisher's fundamental theorem of natural selection to include higher-order cumulants.

#### D. Information associated with conditional selection

Generally, the *mutual information* between two sets of discrete variables, condition $c \in C$ and gene $i \in G$, can be expressed as $I(G;C) = \sum_c \sum_i p(c,i) \ln \frac{p(c,i)}{p(c)p(i)} = \sum_c p(c) \sum_i p(i|c) \ln \frac{p(i|c)}{p(i)} = \sum_c p(c) D_{KL}(p(i|c) \parallel p(i)) = \sum_c p(c) I(G, C = c) = \sum_c p(c) \sum_i p(i|c) I(i; C = c)$. Here, $I(G; C = c)$ can be viewed as the *specific information* for condition $c$, and $I(i; C = c)$ the *pointwise specific information* between a specific gene $i$ and condition $c$ [32, 33]. While $I(G;C)$ and $I(G;C = c)$ are non-negative, $I(i;C = c)$ can take negative values, which indicate that $i$ is misinformative or associates with a loss of information. We can express the information associated with selection at a time $t$ by using the conditional distribution $q_{i,c,t} = q_{i,0} e^{t(\lambda_i + f_{i,c}) - \psi_c}$ and the unperturbed distribution $q_{i,t} = q_{i,0} e^{t\lambda_i - \psi}$, such that

$$I_t(G, C = c) = D_{KL}(q_{i,c,t} \parallel q_{i,t}) = t\langle f_i \rangle_{t,c} - \Delta\psi_c, \quad (9)$$

where $\Delta\psi_c = \ln \frac{\langle e^{t(\lambda_i + f_{i,c})}\rangle_0}{\langle e^{t\lambda_i}\rangle_0}$. Taylor expanding both terms around $t = 0$ gives $t\langle f_{i,c}\rangle_{t,c} = t(\mu_{f_c} + t(\sigma_{f_c}^2 + Cov[\lambda, f_c]_0) + ...)$ and $\Delta\psi_c = t\mu_{f_c} + \frac{t^2}{2}(\sigma_{f_c}^2 + 2Cov[\lambda, f_c]_0) + ...$, which when combined gives

$$I_t(G, C = c) \approx \frac{t^2}{2}\sigma_{f_c}^2, \quad (10)$$

under a second-order approximation. For comparison,

$$\begin{aligned} &D_{KL}(q_{i,0} \parallel q_{i,c,t}) - D_{KL}(q_{i,0} \parallel q_{i,t}) \\ &= -\left\langle \ln \frac{q_{i,c,t}}{q_{i,t}} \right\rangle_0 \approx \frac{t^2}{2}\sigma_{f_c}^2 + t^2 Cov[\lambda, f_c]_0, \end{aligned} \quad (11)$$

which relates the expected conditional divergence to negative selection pressure from the *specific information* plus the covariance between baseline fitness and the interaction effects. The covariance term describes the change in $f_c$ due to selection according to the Price equation [4]. If the baseline fitness and treatment effects are independent, the information associated with a particular condition can be viewed variationally as $\Delta_c D_{KL}(q_{i,0} \parallel q_{i,c,t})$ with respect to a baseline state (i.e., untreated control). Noteworthy, $D_{KL}(q_{i,c,t} \parallel q_{i,t})$ expands in $k$th order coefficients equal to $\frac{1}{(k-2)!k}$ compared to $\frac{1}{k!}$ for $\Delta_c D_{KL}(q_{i,0} \parallel q_{i,c,t})$, representing more sensitivity to higher-order cumulants (Fig. S4C, *lower panel*). Moreover, if the baseline fitness and treatment effects are independent (Fig. S4D), the gene-wise interaction effects can be expressed as the following:

$$\begin{aligned} t\Delta\delta_{i,c} &= t(f_{i,c} - \mu_{f_c}) \\ &= \left[\ln \frac{q_{i,t,c}}{q_{i,t}} - \left\langle \ln \frac{q_{i,t,c}}{q_{i,t}} \right\rangle_0\right] \\ &\approx I_t(i, C = c) + I_t(G, C = c). \end{aligned} \quad (12)$$

Because $I_t(G, C = c) \geq 0$, a zero interaction effect means that the misinformativeness of $i$ (or loss of information in response to mutation $i$) is redundant with the expected genomic information about $c$.

#### E. Simulations

Simulation of selection dynamics was performed using the Gillespie stochastic simulation algorithm [34], with $m$ sub-populations $i$ of competing replicators with initial counts $n_{i,0} \sim \mathcal{P}(n_{init})$, and growth rates $\lambda_i \sim \mathcal{N}(\mu_\lambda, \sigma_\lambda^2)$. To sample $\lambda_i$-values from skewed normal distributions given by $S_\lambda$, we used *rsnorm* implemented in the fGarch package [58]. Each iteration step was executed with a bit of noise $\sim \mathcal{N}(0, \eta)$. The step size for each iteration was given as $\Delta t = -\ln(s_1)/\sum_i^m n_i\lambda_i$, with $s_1 \sim \mathcal{U}(0, 1)$. The replication events were selected for *ith* sub-population corresponding to the smallest integer $k$ satisfying $\sum_{i=1}^k n_i\lambda_i > s_2 \sum_i^m n_i\lambda_i$, with $s_2 \sim \mathcal{U}(0, 1)$. Bottlenecks were introduced by down-sampling the seeding population by a factor $b$ using a multinomial distribution $\mathcal{M}_i(bN, p = n_i/N)$ with $N = \sum_i n_i$, every time $N > n_{init}m$. Recording of the population counts were similarly simulated using $\hat{n}_i \sim \mathcal{M}_i(vn_{init}m, p = n_i/N)$, with $v$ being a subsampling factor. Unless otherwise specified, the simulations were executed with $m = 50$, $n_{init} = 50$, $\eta = 0.05$, $\mu_\lambda = 0.1$, $\sigma_\lambda^2 = 0.2$, $S_\lambda = 0$, $b = 1$, $v = 1$, and run for $max.time = 10$.

#### F. Gene-level fitness perturbation statistics

To assess the significance of a gene deletion causing a fitness perturbation, the estimated fitness perturbations $\delta_j$ from $m$ individual sgRNAs targeting the same gene ($j \in gene_i$) were aggregated into a statistic $\bar{\delta}_{gene_i}$ based on the median, mean, or sum, and evaluated against a $H_0$-distribution for the respective statistic $\bar{\delta}_{0,m}$ generated by randomly sampling the same amount of perturbations from a reference sgRNA control set (non-targeting sgRNA controls, or individual sgRNAs targeting reference non-essential genes). Because any given screen will have genes targeted by a different number of sgRNAs, a $H_0$-distribution was generated for each potential number of sgRNAs. A $H_0$-kernel density estimator $K(\bar{\delta}_{0,m})$ was generated for each distribution, such that under the

$H_0$ the likelihood of observing a genetic fitness perturbation $\bar{\delta}_{gene_i}$ is $p(\bar{\delta}_{gene_i}|K(\bar{\delta}_{0,m}))$. The significance of a gene was evaluated by performing numerical integration of the tail distributions to compute the p-values $Pr(\delta_m \geq \bar{\delta}_{gene}|K(\bar{\delta}_{0,m}))$. To estimate the probability of type I errors, we computed a false discovery rate (FDR) using Bayes' rule:

$$Pr(\bar{\delta}_{0,m}|\bar{\delta}_m \geq \bar{\delta}_{gene}|) = \frac{Pr(\bar{\delta}_m \geq |\bar{\delta}_{gene}|K(\bar{\delta}_{0,m}))Pr(\bar{\delta}_{0,m})}{Pr(\bar{\delta}_m \geq \bar{\delta}_{gene}|K(\bar{\delta}_{gene,m}))}. \tag{13}$$

Here, the sensitivity $Pr(\bar{\delta}_m \geq |\bar{\delta}_{gene,m})$ was estimated from the empirical distributions of $\bar{\delta}_{gene,m}$ for each $m$ number of sgRNAs using the kernel density estimator $K(\bar{\delta}_{gene,m})$. The prior $Pr(\bar{\delta}_{0,m})$ was estimated as the supremum of the cumulative distribution of p-values with respect to a uniform distribution. Density estimators for the $H_0$ and empirical-distributions were generated using the $locfit$ package (v1.5-9.8) in R. We also implemented a local fdr (lfdr) computed from the marginal likelihoods as $Pr(\bar{\delta}_{0,m}|\bar{\delta}_{gene}|) = \frac{p(\bar{\delta}_{gene}|K(\bar{\delta}_{0,m}))Pr(\bar{\delta}_{0,m})}{p(\bar{\delta}_{gene}|K(\bar{\delta}_{gene,m}))}$.

#### G. Alternative single screen analyses

To analyze single CRISPR screens with DESeq2 or Wilcoxon test, we used the *caRpools* package (v0.83, 2016; `https://github.com/boutroslab/caRpools`) [59]. MAGeCK (v0.5.9) was run using Python 3 (v3.11.4) [12].

#### H. CN correction and RCPCz

For ensemble analyses, the variances $\sigma_c^2$ per cell line/condition $c$ were stored for re-scaling the distributions for analyses in Fig. 2A, S5A, and S6A. Before CNc and RCPCz, the $\delta_{gene}$ distributions per cell line were scaled to unit variance and centered around the median.

To correct for CN effects, we used $\log_2(CN_{ratio} + 1)$ data from CCLE (Cancer Cell Line Encyclopedia; DepMap portal 2021), where missing gene or cell line entries were imputed with 1. The CN data were matched with the fitness perturbations, and the random effects model,

$$\delta_{i,c} = \beta_0 + b_{0,i} + (\beta_1 + b_{1,i})x_{i,c} + e_{i,c}, \tag{14}$$

was fitted, with $x_{i,c}$ equal to $\log_2(CN_{ratio}+1)$ for gene $i$ in cell line $c$, the fixed effects $\beta$, and random effects $b \sim N(0, \sigma_b^2)$. The fitness perturbations were subsequently reconstructed with the CN effects subtracted as

$$\delta_{i,c}(CNc) = \beta_0 + b_{0,i} + e_{i,c}. \tag{15}$$

RCPCz was done by standardizing the gene dimension of the fitness perturbation matrices to zero mean and unit variance: $(\delta_{i,j} - \mu_{\delta_i})/\sigma_{\delta_i}$, followed by an SVD and subsequently reconstructing the data without the leading components dominated by batch variation. This reduced gene-level correlations between baseline fitness and interaction effects, while conserving cell-wise skewness (Fig. S4E,F). For Broad and Sanger Institute data individually, the first six components were omitted before reconstructing the datasets and adding back $\mu_{\delta_i}$ and $\sigma_{\delta_i}$. For the Sanger dataset, there were biological replicates for several cells lines, which was either averaged or kept separate. When combined with the Broad dataset, the biological replicates were kept separate. For the combined dataset, we tested the removal of several components (Fig. S10A) to optimize reconstruction. The screens in the combined dataset were first scaled to unit variance and centered around their population medians. To assess the batch variability for each reconstructed data, we computed a mean squared error (MSE) between replicated cell lines across the batches or between different cell lines in the combined ensemble. An optimal removal of 12 components was determined at the inflection point where the fold-change in replication error no longer outpaced the fold-change in between-cell error per component removed (Fig. S10B).

Fitness interactions $\Delta\delta_{i,c}$ for any mutant $i$, were computed by subtracting the mean fitness perturbation $\mu_{\delta_i}$ from $\delta_{i,c}$ values or standardized $\delta_{i,c}(z)$ values across the ensemble of environments $c \in [1, 2, \ldots, R]$.

#### I. Detection of essential genes

Constitutively core essential (CCE) and non-essential (NES) gene sets were retrieved from Hart et al. (2017) [14]. Area under the curves (AUCs) for precision-recall (PRC) and receiver operating characteristic (ROC) curves were computed with the *precrec* package in R. Normalized negative mean difference was computed as $NNMD_j = (\mu_{\delta_{CCE},c} - \mu_{\delta_{NES},c})/\sigma_{\delta_{NES},c}$ per cell line $c$. A high ROC or PRC AUC, while a low NNMD, corresponds to a high detection sensitivity.

#### J. Parametric GO enrichment, semantic similarity clustering and correlation analysis

Enrichment was done on gene ontology (GO) data using the *piano* package in R [40], with "sum" as the gene set statistic, null-hypothesis testing using "geneSampling" with 1000 permutations. Gene set sizes were limited to a range between 5 and 500 genes. Enrichment sensitivity was computed from the number of GO terms with $p \leq 0.01$ divided by the total number of GO terms tested for a given directionality test. PCA was done on standardized $\log_{10}$-transformed p-values from 100 randomly sampled screens.

Enrichment correlation analysis was performed on GO data for biological process (BP), cellular compartment (CC), and molecular function (MF) separately. GO terms were ordered with hierarchical clustering (Euclidean distances and Ward.D2) of semantic similarities based on the Wang method ($1 - SS_{scores}$) [42]. To group the GO terms into domains, we used *cutreeDynamic* from the *dynamicTreeCut* package in R, with $method$ = "*hybrid*", $deepSplit$ = 3, and $minClusterSize = [5, 50, 100, 200, 300]$ for BP, $minClusterSize = [1, 12, 25, 50, 75]$ for CC, and $minClusterSize = [2, 17, 33, 67, 100]$ for MF. The average of $-\log_{10} p$-values was computed per cluster for mixed directional negative and positive tests, and the Pearson correlations between positive and negative clusters statistics were subsequently measured.

#### K. Variable selection with penalized regression

Penalized regression models were fitted using the *glmnet* package in R [60]. Standardized $D_{KL}$ and $D_{KL}/\sigma_\lambda$-values were modeled from $\Delta\delta_i(z)$ using Elastic Net or Lasso regression with different penalty mixture parameters ($\alpha$). Penalties were optimized using MSE and 10-fold cross-validation (CV). For Metascape enrichment [45], variables were selected using Lasso ($\alpha = 1$). Metascape (v3.5.20240101) was run with default settings, but controlling the universe based on the genes present in the analysis.

#### L. Approximation of the MI and $\mathbf{D_{KL}}$ for conditional selection

The following decomposition of the $D_{KL}$ was inspired by the work of Koyama et al. on perturbational thermodynamics [46]. Given marginal and joint probabilities of observing a mutant $i \in G$ and the environmental state $\boldsymbol{\epsilon}$, the *mutual information* between the two variables is defined as $\langle I(G;\boldsymbol{\epsilon})\rangle = \int_{\boldsymbol{\epsilon}} \sum_i p(\boldsymbol{\epsilon}, i) \ln \frac{p(\boldsymbol{\epsilon},i)}{p(\boldsymbol{\epsilon})p(i)} d\boldsymbol{\epsilon} = \int_{\boldsymbol{\epsilon}} p(\boldsymbol{\epsilon}) \sum_i p(i|\boldsymbol{\epsilon}) \ln \frac{p(i|\boldsymbol{\epsilon})}{p(i)} d\boldsymbol{\epsilon}$. Using the replicator equation, $I(G;\boldsymbol{\epsilon})$ can be expressed from the perturbed distribution $q_{i,t\boldsymbol{\epsilon}} = q_{i,0} e^{t(\lambda_i + \boldsymbol{\epsilon}^\top \mathbf{f}_i) - \psi_{\boldsymbol{\epsilon}}}$, and the unperturbed distribution $q_{i,t} = q_{i,0} e^{t\lambda_i - \psi}$:

$$\begin{aligned} \langle I_t(G;\boldsymbol{\epsilon})\rangle &= \langle D_{KL}(q_{i,t,\boldsymbol{\epsilon}} \parallel q_{i,t})\rangle \\ &= \int p(\boldsymbol{\epsilon}) \sum_i q_{i,t\boldsymbol{\epsilon}} \ln \frac{q_{i,t\boldsymbol{\epsilon}}}{q_{i,t}} d\boldsymbol{\epsilon} \\ &= \int p(\boldsymbol{\epsilon}) \sum_i q_{i,t\boldsymbol{\epsilon}} (t\boldsymbol{\epsilon}^\top \mathbf{f}_i - \Delta\psi_{\boldsymbol{\epsilon}}) d\boldsymbol{\epsilon} \\ &= \int p(\boldsymbol{\epsilon}) t\boldsymbol{\epsilon}^\top \langle \mathbf{f}\rangle_{t\boldsymbol{\epsilon}} - \ln \frac{\langle e^{t(\lambda + \boldsymbol{\epsilon}^\top \mathbf{f})}\rangle_0}{\langle e^{t\lambda}\rangle_0} d\boldsymbol{\epsilon} \end{aligned} \tag{16}$$

Identifying $\Delta\psi_{\boldsymbol{\epsilon}}$ as the joint cumulant generating function on the form $K(\boldsymbol{\epsilon}) = \ln M(\boldsymbol{\epsilon})$, we can expand as

$$K(\boldsymbol{\epsilon}) = \sum_r \epsilon_r \frac{\partial K(\boldsymbol{\epsilon})}{\partial \epsilon_r}|_{\epsilon=0} + \frac{1}{2!} \sum_{r,l} \epsilon_r \epsilon_l \frac{\partial^2 K(\boldsymbol{\epsilon})}{\partial \epsilon_r \partial \epsilon_l}|_{\epsilon=0} \cdots, \tag{17}$$

where $\frac{\partial K(\boldsymbol{\epsilon})}{\partial \epsilon_r} = \frac{1}{M(\boldsymbol{\epsilon})} \frac{\partial M(\boldsymbol{\epsilon})}{\partial \epsilon_r} = t\langle f_r\rangle_{t\boldsymbol{\epsilon}}$ and $\frac{\partial K(\boldsymbol{\epsilon})}{\partial \epsilon_r \partial \epsilon_l} = \frac{\partial}{\partial \epsilon_l}(\frac{1}{M(\boldsymbol{\epsilon})} \frac{\partial M(\boldsymbol{\epsilon})}{\partial \epsilon_r}) = \frac{\partial \langle f_r\rangle_{t\boldsymbol{\epsilon}}}{\partial \epsilon_l} = t^2(\langle f_r f_l\rangle_{t\boldsymbol{\epsilon}} - \langle f_r\rangle_{t\boldsymbol{\epsilon}} \langle f_l\rangle_{t\boldsymbol{\epsilon}})$. Similarly, we can Taylor expand $\langle \mathbf{f}\rangle_{t\boldsymbol{\epsilon}}$, such that

$$\begin{aligned} t\boldsymbol{\epsilon}^\top \langle \mathbf{f}\rangle_{t\boldsymbol{\epsilon}} &= t\sum_r \epsilon_r \langle f_r\rangle_t + t\sum_{r,l} \epsilon_r \epsilon_l \frac{\partial \langle f_r\rangle_{t\boldsymbol{\epsilon}}}{\partial \epsilon_l}|_{\epsilon=0} \cdots. \\ &= \sum_r \epsilon_r \frac{\partial K(\boldsymbol{\epsilon})}{\partial \epsilon_r}|_{\epsilon=0} + \sum_{r,l} \epsilon_r \epsilon_l \frac{\partial^2 K(\boldsymbol{\epsilon})}{\partial \epsilon_r \partial \epsilon_l}|_{\epsilon=0} \cdots. \end{aligned} \tag{18}$$

Thus, combining these results, we get

$$\begin{aligned} t\boldsymbol{\epsilon}^\top \langle \mathbf{f}\rangle_{t\boldsymbol{\epsilon}} - \Delta\psi_{\boldsymbol{\epsilon}} &= \frac{t^2}{2} \sum_{r,l} \epsilon_r \epsilon_l (\langle f_r f_l\rangle_t - \langle f_r\rangle_t \langle f_l\rangle_t) \cdots \\ &= \frac{t^2}{2} \boldsymbol{\epsilon}^\top \langle (\mathbf{f} - \langle \mathbf{f}\rangle_t)(\mathbf{f} - \langle \mathbf{f}\rangle_t)^\top \rangle_t \boldsymbol{\epsilon} \cdots \\ &= \frac{t^2}{2} \boldsymbol{\epsilon}^\top \mathbf{C_t} \boldsymbol{\epsilon} \cdots, \end{aligned} \tag{19}$$

with $k$th-order coefficients equal to $\frac{1}{(k-2)!k}$, and the first term corresponding to the second-order approximation, which gives $I_t(G;\boldsymbol{\epsilon}) \approx \frac{t^2}{2} \int p(\boldsymbol{\epsilon}) \boldsymbol{\epsilon}^\top \mathbf{C_t} \boldsymbol{\epsilon} d\boldsymbol{\epsilon}$. Equivalently, if we expand $\langle \mathbf{f}\rangle_{t\boldsymbol{\epsilon}}$ and $\Delta\psi_{\boldsymbol{\epsilon}}$ around $t = 0$, each second-order term will include the additional covariance $t^2 \boldsymbol{\epsilon}^\top \langle (\mathbf{f} - \langle \mathbf{f}\rangle_0)(\lambda - \mu_\lambda)^\top \rangle_0$, which will cancel each other out when combined. Moreover, assuming no conditional selection prior to the initialization of an experiment, $\lambda$ and $\mathbf{f}$ should be independent. Thus, it follows that under a second-order approximation $I_t(G;\boldsymbol{\epsilon}) \approx \frac{t^2}{2} \int p(\boldsymbol{\epsilon}) \boldsymbol{\epsilon}^\top \mathbf{C_0} \boldsymbol{\epsilon} d\boldsymbol{\epsilon}$.

We can analyze the change in $D_{KL}(q_{i,t,\boldsymbol{\epsilon}} \parallel q_{i,t}) = t\boldsymbol{\epsilon}^\top \langle \mathbf{f}\rangle_{t\boldsymbol{\epsilon}} - \Delta\psi_{\boldsymbol{\epsilon}}$ as a function of the environment by computing its gradient with respect to $\boldsymbol{\epsilon}$:

$$\begin{aligned} \nabla_{\boldsymbol{\epsilon}} D_{KL}(q_{i,t,\boldsymbol{\epsilon}} \parallel q_{i,t}) &= t\langle \mathbf{f}\rangle_{t\boldsymbol{\epsilon}} + t\boldsymbol{\epsilon}^\top \nabla_{\boldsymbol{\epsilon}} \langle \mathbf{f}\rangle_{t\boldsymbol{\epsilon}} - t\langle \mathbf{f}\rangle_{t\boldsymbol{\epsilon}} \\ &= t\boldsymbol{\epsilon}^\top \nabla_{\boldsymbol{\epsilon}} \langle \mathbf{f}\rangle_{t\boldsymbol{\epsilon}}. \end{aligned} \tag{20}$$

This expression can be approximated as

$$\boldsymbol{\epsilon}^\top \nabla_{\boldsymbol{\epsilon}} \langle \mathbf{f}\rangle_{t\boldsymbol{\epsilon}} \approx \langle \mathbf{f}\rangle_{t\boldsymbol{\epsilon}} - \langle \mathbf{f}\rangle_0 \approx t\mathbf{C_0}\boldsymbol{\epsilon} \tag{21}$$

in a linear response regime and given the second-order approximation. This perturbation can be recognized as an upper bound for the $D_{KL}$ via Jensen's inequality [46],

$$t\boldsymbol{\epsilon}^\top \langle \mathbf{f}\rangle_{t\boldsymbol{\epsilon}} - \Delta\psi_{\boldsymbol{\epsilon}} \le t\boldsymbol{\epsilon}^\top (\langle \mathbf{f}\rangle_{t\boldsymbol{\epsilon}} - \langle \mathbf{f}\rangle_0). \tag{22}$$

Finally, by computing

$$\begin{aligned}\langle \ln \frac{q_{i,t\epsilon}}{q_{i,t}} \rangle_0 &= t\epsilon^\top \langle \mathbf{f} \rangle_0 - \Delta\psi_\epsilon \\ &\approx -\frac{t^2}{2}\epsilon^\top \mathbf{C_0}\epsilon - t^2\epsilon^\top \langle (\mathbf{f} - \langle \mathbf{f} \rangle_0)(\lambda - \mu_\lambda)^\top \rangle_0,\end{aligned} \tag{23}$$

we can set

$$\begin{aligned}\langle \ln \frac{q_{i,t\epsilon}}{q_{i,t}} \rangle_{t\epsilon} - \langle \ln \frac{q_{i,t\epsilon}}{q_{i,t}} \rangle_0 &= t\epsilon^\top (\langle \mathbf{f} \rangle_{t\epsilon} - \langle \mathbf{f} \rangle_0) \\ &\approx t^2\epsilon^\top \mathbf{C_0}\epsilon + t^2\epsilon^\top \langle (\mathbf{f} - \langle \mathbf{f} \rangle_0)(\lambda - \mu_\lambda)^\top \rangle_0 \\ &= t^2\epsilon^\top \mathbf{C_0}\epsilon,\end{aligned} \tag{24}$$

given the second-order approximation and independence between $\lambda$ and $\mathbf{f}$. These relations allow us to approximate the environmental perturbations based on the expected change in $\mathbf{f}$ as $t\epsilon \approx \mathbf{C_0}^{-1}(\langle \mathbf{f} \rangle_{t\epsilon} - \langle \mathbf{f} \rangle_0)$.

Consequently, the $D_{KL}$ can be expressed as an expectation of $\mathbf{f}$ as:

$$\begin{aligned}\frac{t^2}{2}\epsilon^\top \mathbf{C_0}\epsilon &\approx \frac{1}{2}(\langle \mathbf{f} \rangle_{t\epsilon} - \langle \mathbf{f} \rangle_0)^\top \mathbf{C_0}^{-1}(\langle \mathbf{f} \rangle_{t\epsilon} - \langle \mathbf{f} \rangle_0) \\ &= \frac{1}{2}(\langle \mathbf{f} \rangle_{t\epsilon} - \langle \mathbf{f} \rangle_0)^\top \mathbf{V}\mathbf{\Sigma}^{-2}\mathbf{V}^\top (\langle \mathbf{f} \rangle_{t\epsilon} - \langle \mathbf{f} \rangle_0) \\ &\approx \frac{N-1}{2}(\langle \mathbf{u} \rangle_{t\epsilon_*}^\top \mathbf{\Sigma}\mathbf{V}^\top)\mathbf{V}\mathbf{\Sigma}^{-2}\mathbf{V}^\top (\langle \mathbf{u} \rangle_{t\epsilon_*}^\top \mathbf{\Sigma}\mathbf{V}^\top)^\top \\ &= \frac{N-1}{2}\langle \mathbf{u} \rangle_{t\epsilon_*}^\top \langle \mathbf{u} \rangle_{t\epsilon_*},\end{aligned} \tag{25}$$

with $(\mathbf{f}_i - \langle \mathbf{f} \rangle_0)^\top = \sqrt{N-1}\mathbf{u}_i^\top \mathbf{\Sigma}\mathbf{V}^\top$ from the SVD: $\Delta\mathbf{F} = \mathbf{U}\mathbf{D}\mathbf{V}^\top$, with singular values $\mathbf{D}$, which are related to the diagonalized variance as $\mathbf{\Sigma}^2 = \frac{1}{N-1}\mathbf{D}^2$.

Assuming independence between $\lambda$ and $\mathbf{f}$, the conditional selection of a mutant $i$ in environment $\epsilon$ can be expressed by the *pointwise mutual information*, $I(i;\epsilon)$, as:

$$\begin{aligned}\ln \frac{q_{i,t\epsilon}}{q_{i,t}} &= t\epsilon^\top \mathbf{f}_i - \Delta\psi_\epsilon \\ &\approx t\epsilon^\top (\mathbf{f}_i - \langle \mathbf{f} \rangle_0) - \frac{t^2}{2}\epsilon^\top \mathbf{C_0}\epsilon \\ &\approx (\langle \mathbf{f} \rangle_{t\epsilon} - \langle \mathbf{f} \rangle_0)^\top \mathbf{C_t}^{-1}(\mathbf{f}_i - \langle \mathbf{f} \rangle_0) \\ &\quad - \frac{1}{2}(\langle \mathbf{f} \rangle_{t\epsilon} - \langle \mathbf{f} \rangle_0)^\top \mathbf{C_t}^{-1}(\langle \mathbf{f} \rangle_{t\epsilon} - \langle \mathbf{f} \rangle_0) \\ &\approx (N-1)[\langle \mathbf{u} \rangle_{t\epsilon_*}^\top \mathbf{u}_i - \frac{1}{2}\langle \mathbf{u} \rangle_{t\epsilon_*}^\top \langle \mathbf{u} \rangle_{t\epsilon_*}].\end{aligned} \tag{26}$$

Maximum conditional selection of mutant $i$ involves finding $\epsilon_*$ such that:

$$I(i;\epsilon)_{max} = \max_{\substack{\epsilon_* \\ \langle \mathbf{u} \rangle_{t\epsilon_*} = \mathbf{u}_i}} \ln \frac{q_{i,t\epsilon}}{q_{i,t}} \approx \frac{N-1}{2}\mathbf{u}_i^\top \mathbf{u}_i, \tag{27}$$

which represents the maximum amount of environmental information encoded in $i$.

### M. Estimating the MI between mutants

Because we can assume independent growth, the joint probability of observing any two mutants $i$ and $j$ is conditionally independent given $\epsilon$, such that $P(i,j) = \int p(\epsilon)P(i|\epsilon)P(j|\epsilon)d\epsilon$. Thus, the pointwise mutual information $I(i;j) = \ln \frac{P(i,j)}{P(i)P(j)}$ can be expressed as a function of growth perturbations as follows:

$$I(i;j) = \ln \int p(\epsilon)\frac{q_{i,t\epsilon}q_{j,t\epsilon}}{q_{i,t}q_{j,t}}d\epsilon \geq \int p(\epsilon)\ln \frac{q_{i,t\epsilon}q_{j,t\epsilon}}{q_{i,t}q_{j,t}}d\epsilon, \tag{28}$$

with the latter step given by Jensen's inequality. Plugging in equation 13, we get $I(i;j) \geq (N-1)\int p_{\epsilon_*}\langle \mathbf{u} \rangle_{t\epsilon_*}^\top \mathbf{u}_i + \langle \mathbf{u} \rangle_{t\epsilon_*}^\top \mathbf{u}_j - \langle \mathbf{u} \rangle_{t\epsilon_*}^\top \langle \mathbf{u} \rangle_{t\epsilon_*} d\epsilon_*$. In a similar fashion, the maximum co-varying selection of $i$ and $j$ involves finding $\epsilon_*$ such that:

$$\begin{aligned}I(i;j)_{max} &= \max_{\substack{\epsilon_* \\ \langle \mathbf{u} \rangle_{t\epsilon_*} = \mathbf{u}_i \vee \langle \mathbf{u} \rangle_{t\epsilon_*} = \mathbf{u}_j}} \ln \frac{q_{i,t\epsilon}q_{j,t\epsilon}}{q_{i,t}q_{j,t}} \\ &\approx (N-1)\mathbf{u}_i^\top \mathbf{u}_j,\end{aligned} \tag{29}$$

which represents the maximum mutual dependence between $i$ and $j$, or the maximum amount of information one genetic trait encodes about the selection of another. Notably, if we compute the mutual information between a gene $i$ and itself, we get $I(i;i)_{max} \approx (N-1)\mathbf{u}_i^\top \mathbf{u}_j$, which is referred to as the maximum self-information. We can see that this value is related to $I(i;\epsilon)_{max}$ and underscores the squared norm of normalized projection coordinates as an estimator for the maximally encoded information per gene. The normalized mutual information was computed as $\cos(\mathbf{u}_i, \mathbf{u}_j) = \frac{\mathbf{u}_i^\top \mathbf{u}_j}{||\mathbf{u}_i|| ||\mathbf{u}_j||}$, where $||\mathbf{u}_i|| = \sqrt{\mathbf{u}_i^\top \mathbf{u}_i}$.

### N. Estimating MI from fitness perturbations

With linear combinations in the response coefficients $\mathbf{f}_i$ determining the perturbations in fitness $\lambda_{i,\epsilon}$, and the normalized projection coordinates $\mathbf{u}_i$ characterizing the independent components contributing to the fitness response, we can set the following:

$$\begin{aligned}\Delta\delta_{i,\epsilon} = \delta_{i,\epsilon} - \delta_i &= (\lambda_{i,\epsilon} - \mu_{\lambda_\epsilon}) - (\lambda_i - \mu_\lambda) \\ &= (\mathbf{f}_i - \langle \mathbf{f} \rangle_0)^\top \epsilon \\ &= \mathbf{u}_i^\top \mathbf{D}\mathbf{V}^\top \epsilon \\ &= \mathbf{u}_i^\top \mathbf{D}_* \mathbf{w}_\epsilon,\end{aligned} \tag{30}$$

where $\mathbf{D}_*$ is composed of diagonalized singular values related to the standard deviations in fitness interactions, $\frac{1}{\sqrt{N-1}}\mathbf{D}_* = \mathbf{\Sigma}_* = diag[\sigma_{\delta,1}, \sigma_{\delta,2}, ..., \sigma_{\delta,R}]$. Moreover, $\mathbf{w}_\epsilon$ is a left singular vector containing the decomposed perturbations corresponding to a specific environment $c$ with perturbations $\epsilon$. Thus, the mutual information ($MI$) content under maximum selection can be retrieved from the projection matrix $\mathbf{U}\mathbf{U}^\top$ based on the left singular vectors $\mathbf{U}$ of the SVD $\mathbf{\Delta} = \mathbf{U}\mathbf{D}_*\mathbf{W}^\top$, which effectively executes a PCA on the fitness matrix $[\boldsymbol{\delta}_1, \boldsymbol{\delta}_2, ..., \boldsymbol{\delta}_N]$, where the mean-subtracted matrix $\mathbf{\Delta} = [\Delta\boldsymbol{\delta}_1, \Delta\boldsymbol{\delta}_2, ..., \Delta\boldsymbol{\delta}_N]^\top$ is the fitness interaction matrix. The normalized mutual information ($MI_{norm}$) is equivalent to computing it from the Pearson correlation coefficients between the rows in $\mathbf{U}$ ($PCC_U$). For comparison, we also computed the correlations between the principal components $\mathbf{G}_* = \mathbf{U}\mathbf{\Sigma}_*$ ($PCC_{PC}$) and fitness perturbations $[\boldsymbol{\delta}_1, \boldsymbol{\delta}_2, ..., \boldsymbol{\delta}_N]$ ($PCC_{fitness}$).

#### O. Differential MI in cell-type subspaces

The rows of $\mathbf{W}$ belonging to a cell-type subset $S$ were removed, and the projection matrix was then rotated using $\mathbf{W}_{\setminus S}^\top \mathbf{W}_{\setminus S}$, with $I(i; \boldsymbol{\epsilon}_{\setminus S})_{max} \sim (\mathbf{u}_i^\top \mathbf{u}_i)_{\setminus S} = \mathbf{u}_i^\top \mathbf{W}_{\setminus S}^\top \mathbf{W}_{\setminus S} \mathbf{u}_i$, corresponding to the information gain under maximum selection within the subspace excluding cell lines $c \in S$. The normalized mutual information for maximal co-varying selection of two mutants was similarly computed as $\cos(\mathbf{u}_i, \mathbf{u}_j)_{\setminus S} = \frac{(\mathbf{u}_i^\top \mathbf{u}_j)_{\setminus S}}{||\mathbf{u}_i||_{\setminus S}||\mathbf{u}_j||_{\setminus S}}$.

To retrieve the maximum mutual dependence between genetic traits specific to a given cell type, we computed the differential normalized mutual information $\Delta MI_{norm}$ as $\Delta_S \cos(\mathbf{u}_i, \mathbf{u}_j) = \cos(\mathbf{u}_i^\top \mathbf{u}_j) - \cos(\mathbf{u}_i^\top \mathbf{u}_j)_{\setminus S}$. To retrieve cell-type-specific information gain under maximum selection, we computed the differential mutual information $\Delta MI$ between the native and rotated projection matrix, normalized on the respective matrix ranks, $\Delta_S I(i; \boldsymbol{\epsilon})_{max} = I(i; \boldsymbol{\epsilon})_{max}/R - I(i; \boldsymbol{\epsilon}_{\setminus S})_{max}/R_{\setminus S}$. For comparison we computed differential mean fitenss perturbation per gene $\Delta_S \mu_{\delta_i} = \mu_{\delta_i} - \mu_{\delta_i, \setminus S}$ and the differential fitness variance per gene $\Delta_S \sigma^2_{\delta_i} = \sigma^2_{\delta_i} - \sigma^2_{\delta_i, \setminus S}$.

To generate the cell-type subsets, the following queries were used on the respective columns within the DepMap sample info data file (DepMap portal 2021): *liver*, *kidney* and *skin* from lineage; *t_cell* and *b_cell* from lineage_sub_subtype; *glioma* from lineage_subtype for glia; *neuroblastoma* and *medulloblastoma* from lineage_subtype and *autonomic_ganglia* from sample_collection_site for neurons. Similarly, for age group subsets we sampled conditions based on the reported donor age of the screened cells.

For predicting donor age based on the differential scores per corresponding cell environment $c$, the sample-specific $\Delta_S I(i; \boldsymbol{\epsilon}_{\setminus c})_{max}$ values were estimated as: $\mathbf{u}_i^\top \mathbf{u}_i / R - \mathbf{u}_i^\top \mathbf{w}_c^\top \mathbf{w}_c \mathbf{u}_i$. Similarly, this was compared with differential fitness (i.e. fitness interactions), $\Delta\delta_{i,c} = \delta_{i,c} - \mu_{\delta_i}$, and sample-specific fitness variance $(\Delta\delta_{i,c})^2$. These values were fitted to predict the log-transformed donor age $\log_{10}(age_c + 1)$ using penalized regression with the *glmnet* package in R [60].

#### P. FLEX and STRING enrichment analysis

Precision-recall of CORUM complexes was performed with the *FLEX* package (`https://github.com/csbio/FLEX_R`) [21].

STRING enrichments were based on the STRING v10 database [61] and performed on all detected interactions or protein-protein interactions (PPIs) with $experiments\_score \geq 0$ or $experiments\_transferred\_score \geq 0$. Genome-wide enrichments were computed by finding the frequency of true interactions per similarity rank and dividing this by a background frequency computed from the average of 20 iterations randomly sampling gene pairs from the entire similarity matrix. For cumulative enrichments the frequency up to a similarity rank was computed.

#### Q. UMAP

UMAP was performed using the *umap* package (v0.2.9.0) in R [48]. Data implementation and distance metrics were defined as indicated, and all embeddings were performed with $n\_neighbors = 25$. For optimization of the embeddings, $n\_epochs = 1000$, and for final embeddings $n\_epochs = 3000$.

#### R. GSEA and Fisher's exact test

GSEA and over-representation analyses using Fisher's exact test were performed using the *clusterProfiler* package in R [47]. For k-means clustering analysis of UMAP GO-BP enrichment, the maximum adjusted $p$-value was retrieved per unique GO term before computing the average $-\log_{10} p_{adj}$-value or the number of GO terms with $p_{adj} \leq 0.01$. When comparing different fitness computation methods, only the intersecting genes between all approaches were embedded and analyzed. Grouping of GO terms relevant for age groups was done using the following regular expression search queries: *development*, *differentiation*, and *growth*, respectively; *immun|inflamm|antigen|mhc* for *immunity*; *ubiquitin|proteasome|lysosom|* and *phagy|autophagosome|atg|phagophore* for *proteostasis*.

#### S. Network analyses and SAFE

Assuming sparsity of true interactions, networks were generated for different similarity metrics by randomly sampling 100000 gene-pairs and setting a quantile cutoff based on the absolute similarity score. Genome-wide STRING-PPI enrichments per network were computed as a classical gene-wise over-representation of edge interaction frequency over the total interaction frequency per gene: $\log_2\left(\frac{k_{PPI,i} \times n_i}{k_i \times n_{PPI,i}}\right)$, where $k$ indicates the number of edges (node degree) and $n$ indicates the total pairs for gene $i$. For all genes with $k_{PPI,i} > 0$, the enrichment scores were summed to find an optimal cutoff ($p \leq 0.001$).

$\Delta MI_{norm}$-based cell-type networks were generated in a similar way by finding edges above a cutoff ($p \leq 0.001$) for both the differential score $\Delta_S \cos(\mathbf{u}_i, \mathbf{u}_j)$ and adjusted similarity score $\cos(\mathbf{u}_i, \mathbf{u}_j) + \Delta_S \cos(\mathbf{u}_i, \mathbf{u}_j)$. For the analysis of age-group-specific network structures, we only set a cutoff based on the differential score.

SAFE (Spatial analysis and functional enrichment) was performed using Cytoscape (v3.9.1; Java 11.0.6) [49]. Networks were embedded using an unweighted, spring-embedded layout algorithm. Enrichments were performed with $distance = map\_weighted$, $threshold = 0.75$, and composite maps were generated using $min.\_landscape\_size = 5$, $similarity = Jaccard$, and $threshold = 0.75$. For genome-wide networks, the background was set as the nodes in the networks. For cell-type networks, the background was set as the nodes in the attribute file, and multi-regional landscapes were included.

Network structures were checked for power-law distributions by measuring the log-linearity between the node degree $k$ and degree density $P(k)$. Hub genes for $MI$-based networks were defined as the top 5% quantile. To score the pleiotropy, we computed the variance in Euclidean distance for all the edges of a node in the UMAP: $\sigma^2_{d_{UMAP}}$, where $d_{UMAP_{i,j}} = \sqrt{(x_i - x_j)^2 + (y_i - y_j)^2}$ is the distance between gene $i$ and $j$. Significant changes in pleiotropy genes were identified using a chi-square test for the difference in $\sigma^2_{d_{UMAP},i}$ for gene $i$ and the average $\sigma^2_{d_{UMAP}}$ in the network, with $k-2$ degrees of freedom. The $p$-values were adjusted using the Benjamini-Hochberg procedure, and low- and high-pleiotropy genes were selected based on $p_{adj} \leq 0.05$. Significant differences in mean fitness $\delta$ or $MI$ among gene hubs were assessed using a Wilcoxon rank sum test.

### T. Public CRISPR screen datasets

For the analysis of single CRISPR screens, we used publicly available data from Hart et al. (2015) and Wang et al. (2015)[9, 62].

For ensemble analysis, the datasets from the Sanger Institute, including raw readcounts, log-fold changes and corrected log-fold changes using CRISPRcleanR (CCR), were retrieved from the Project Score portal (`https://score.depmap.sanger.ac.uk/`) [36]. The Achilles datasets from the Broad Institute, including raw readcounts, guide map, and replicate map, were retrieved from the DepMap portal (`https://depmap.org/portal/download/all/`) [15], under the 20Q3 release. Similarly, CERES fitness scores, with and without RCPC, were retrieved from the 20Q3 release [19]. Chronos fitness scores were retrieved from the 21Q2 release [17]. The CERES data, combining Achilles and Project Score with batch correction using Harmonia, were retrieved from the 21Q2 release [22].

For equal comparison of functional similarity scores with those from Wainberg et al. we also performed the FPA on Broad data from the 18Q3 DepMap release. The Wainberg network data were downloaded from their respective GitHub repository (`https://github.com/kundajelab/coessentiality`) [20].

### U. Drug-perturbed CRISPR screen

#### 1. *Production of lentiviral vectors*

The single-vector lentiviral GeCKO system of mouse GeCKO library v2 (library A and B) was obtained from Addgene. The mouse GeCKO library v2 target 20,661 protein-coding genes with 6 independent sgRNAs, as well as 1175 miRNA-coding regions with 4 independent sgRNAs. DNA was amplified according to the manufacturer's instructions. Lentiviral vectors production: HEK293T were plated in T175 flasks in IMEM medium containing 10% FBS and Pennicillin/Streptomycin. When cells were 80% confluent, the medium was changed (15 ml), and transfection performed using Polyethylenimine (PEI) (Polysciences 23966), pH 4.5, with ratio PEI:DNA = 6:1. For one flask 5 µg of VSVG plasmid (Addgene 12259) + 7.5 µg of REV plasmid (Addgene 12253) + 7.5 µg of RRE plasmid (Addgene 12251) + 1 µg of GeCKO vectors libraryA + 1 µg of GeCKO vectors libraryB (=22 µg DNA) in serum free IMEM was added to 132 µl of PEI pH 4.5 (1 mg/ml), vortexed for 10 seconds and left for 7 minutes at room temperature, after which the PEI-DNA mix was added to HEK293T cells. 24h later, medium was changed (10 ml) and sodium butyrate (Aldrich 303410) was added to reach a final concentration of 5 mM; medium was changed again 8 hours after the addition of sodium butyrate. Supernatants were collected 48h and 72h after transfection, and filtered using a 0.2 µm filter. Lentiviral vectors were concentrated by centrifugation at 40,000 g for 90 min in 50 ml Polybrene tubes, pellets were resuspended overnight in PBS at 4°C, aliquoted and stored at -80°C.

#### 2. *Transduction and pooled CRISPR/Cas9 screen*

35 million BaF3-BCR-ABL1-T315I cells at a concentration 1 million cells in 480 µl medium containing HEPES + polybrene at a final concentration of 5 µg/ml were plated in 24-well plates (1 million cells per well). After 30 minutes, 20 µl of lentiviral vectors were added in each well with an approximate multiplicity of infection of 0.25. After 3 hours, 1 ml medium was added to each well. Twelve hours later, cells from three wells were pooled and transferred to T175 flasks with 20 ml of medium. 48h after transduction, cells were treated with puromycin at a final concentration of 0.85 µg/ml. The next days, cells were centrifuged at 900 RPM for 5 min

and resuspended in fresh medium to remove the dead cells and plated in 15 cm dishes. Cells were kept under puromycin selection until 11 days after transduction. To passage cells, cells were pooled, counted, and plated in 35 dishes (15 cm) containing 2 million cells each in 20 ml medium + puromycin. 11 days after transduction, 70 millions cells were harvested (experimental T0), and the rest of cells were separated into two groups: one group received ponatinib treatment and one group received DMSO treatment. For each group, 28 dishes (15 cm) were plated with 2 million cells in 20 ml medium in each dish. Every 2 days (for a total time of 14 days), 70 million cells were harvested and the rest were passaged into fresh medium containing either ponatinib or DMSO (0.133%). The initial concentration of ponatinib (from time 0 to day 2) was 1 nM, but we observed that this concentration did not confer a significant fitness defect to the cells. From day 2 to day 4, the ponatinib concentration was increased to 7 nM, after which 60% of cells were killed compared to the DMSO control. From day 4 to day 6, the ponatinib concentration was slightly decreased to 5 nM resulting in approximately 50% killing of cells relative to DMSO. The ponatinib concentration was then kept at 5 nM and the pool of ponatinib-treated cells progressively became resistant to the treatment. To passage cells during the 14-day experiment, for each condition cells were mixed, counted, and 56 million cells were put in 560 ml medium, ponatinib or DMSO was added and cells were plated in 28 dishes (15 cm dish, 20 ml of cell in each). Harvesting cells: 70 million cells were centrifuged for 5 min at 1500 rpm, washed twice in PBS and the pellet was frozen. In the end, the samples that were harvested and sequenced were T0, and ponatinib and DMSO treatments after 2, 4, 10 and 14 days. We harvested two samples of cells treated with ponatinib for 14 days, to determine technical reproducibility of our method.

#### 3. *DNA preparation and sequencing*

DNA was extracted using QiaAmp DNA Blood Maxi Kit (51192). A first PCR round (25 cycles) was then performed on the extracted genomic DNA from all collected samples. The primers used were Gecko_v2Adaptor_F and Gecko_v2Adaptor_R.For each sample, 24 reactions with 3µg of DNA in each was performed with Herculase II fusion DNA polymerase (Agilent 600677,600679) following the manufacturer's protocol with addition of DMSO to a final concentration of 2%. PCR products were purified using agarose gel purification, and a second-round PCR (8 cycles with Herculase II) was performed using the following primers for each sample: T0: F02+R02; DMSO 4 days: F04+R04; Ponatinib 4 days: F05+R05; DMSO 10 days: F07+R07; Ponatinib 10 days: F08+R08; DMSO 14 days: F09+R09; Ponatinib 14 days: F01+R01; Ponatinib 14 days (2nd sample): F06+R06. For the second-round PCR, the DMSO 2 days sample was amplified twice with different primers (as a control), and these two samples were referred to as 'DMSO 2 days-1' and 'DMSO 2 days-2', using primer combinations F03+R03 and F11+R11, respectively.All PCR products were purified using agarose gel purification prior to Illumina sequencing.

### V. Software

All analyses, unless otherwise specified, were run using R (v4.2.1). The FPA toolbox for analysis of single and ensemble CRISPR screens is publicly available as an R package *PANIC* (Perturbational Analysis of CRISPR screens).

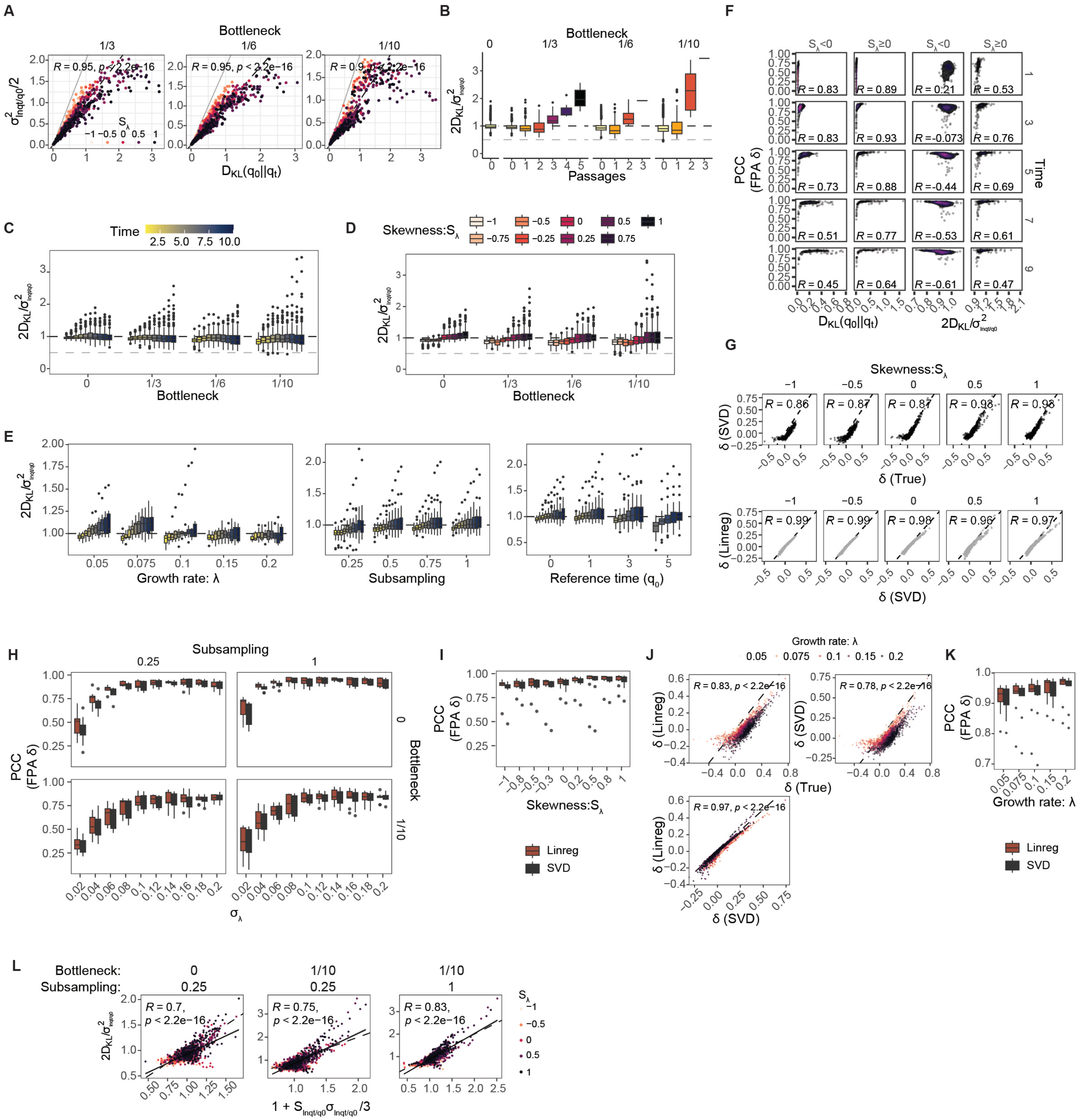


**Fig. S1. Stochastic simulation results.** **A** $D_{KL}$ as a function of empirical variance for different fitness skewness thresholds and bottlenecks. Increasing the down-sampling factor ($b$, indicated *top*) introduces more noise. **B** Relative divergences across passage numbers, representing the number of bottlenecks a population undergoes, for varying down-sampling factors. **C** Relative divergence over time for different bottlenecks. **D** Relative divergence per fitness skewness threshold for different bottlenecks. **E** Relative divergence over time for different population growth rates, subsampling factors ($v$), and reference time points ($t_0$). **F** Pearson correlations for FPA-estimated fitness at different time points relative to $D_{KL}$ and relative divergence for positive and negative fitness skewness. $R$ indicates the Spearman correlation. **G** Comparison between SVD-estimated fitness and true fitness (*top panels*), and between SVD- and regression-estimated fitness (*bottom panels*) for various fitness skewness thresholds. $R$ indicates the Pearson correlation. **H** Pearson correlations between FPA-estimated and true fitness across fitness variance thresholds, bottlenecks ($b$, indicated *right*), and subsampling factors ($v$, indicated *top*). **I** Pearson correlations between FPA-estimated and true fitness for different fitness skewness thresholds. **J** Influence of population growth rate on fitness estimation using linear regression and SVD. $R$ indicates the Pearson correlation. **K** Pearson correlations between FPA-estimated and true fitness across population growth rates. **L** Relative divergence estimated from empirical skewness for various bottlenecks and subsampling factors.

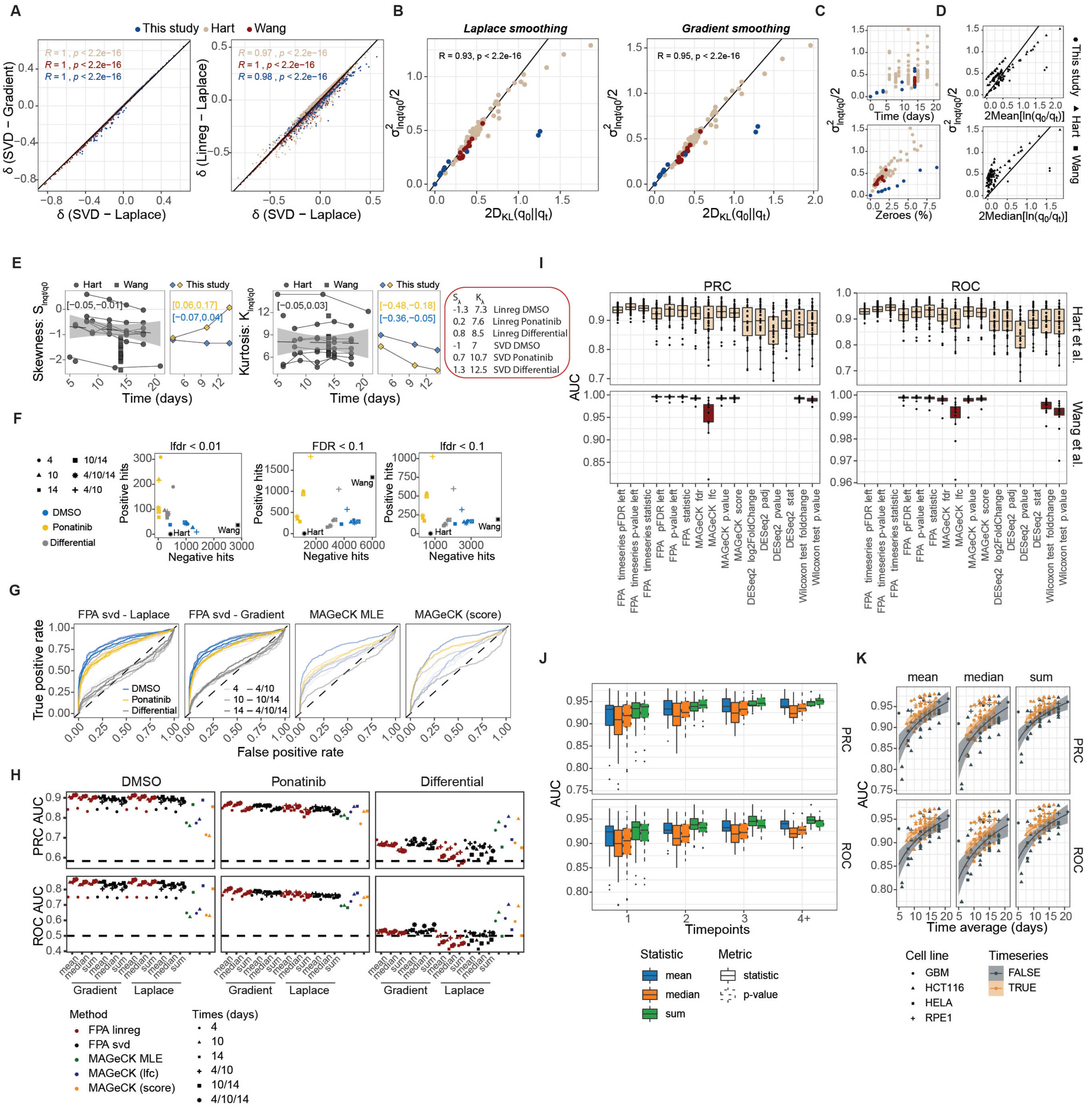

**Fig. S2. FPA benchmarking of individual CRISPR screens.** **A** Comparison of fitness estimation methods. The *left panel* compares different smoothing techniques, while the *right panel* compares SVD and linear regression. $R$ indicates the Pearson correlation. **B** $D_{KL}$ as a function of empirical variance for various smoothing techniques. $R$ indicates the Pearson correlation. **C** Changes in empirical variance across different sample time points (*top*) and percentages of zero reads (*bottom*). **D** Empirical variance compared to the mean (*top*) and median (*bottom*). **E** Empirical skewness (*left panels*) and kurtosis (*right panels*) over time. Values in brackets represent the 95% CI for the slope. The *right box* shows estimated fitness skewness and kurtosis for time series analysis of the Ponatinib response screen using linear regression or SVD. **F** Shifts in the number of significant positive and negative perturbations in response to selective pressure, using different FDR and lfdr cutoffs. Black dots represent the average from other CRISPR screens. **G** ROC curves for detecting essential genes using SVD FPA with various smoothing techniques and mean as the gene-level statistic, compared to MAGeCK using RRA scores or MLE beta values. **H** PRC and ROC AUC summary results for various FPA configurations and MAGeCK outputs. **I** PRC and ROC AUC summary results for FPA, MAGeCK RRA, DESeq2, or Wilcoxon test statistics and significance scores. FPA was performed with linear regression and sum as the gene-level statistic, the best-performing procedure for the Hart et al. dataset. For time series analysis, all possible time combinations were tested. **J** PRC and ROC AUC for various FPA gene-level statistics and significance scores as a function of increasing time points in the time series analysis. **K** PRC and ROC AUC for various time averages during time series FPA using linear regression. Results for different gene-level statistics are shown. Panels J and K represent analyses of time series CRISPR screens from Hart et al.

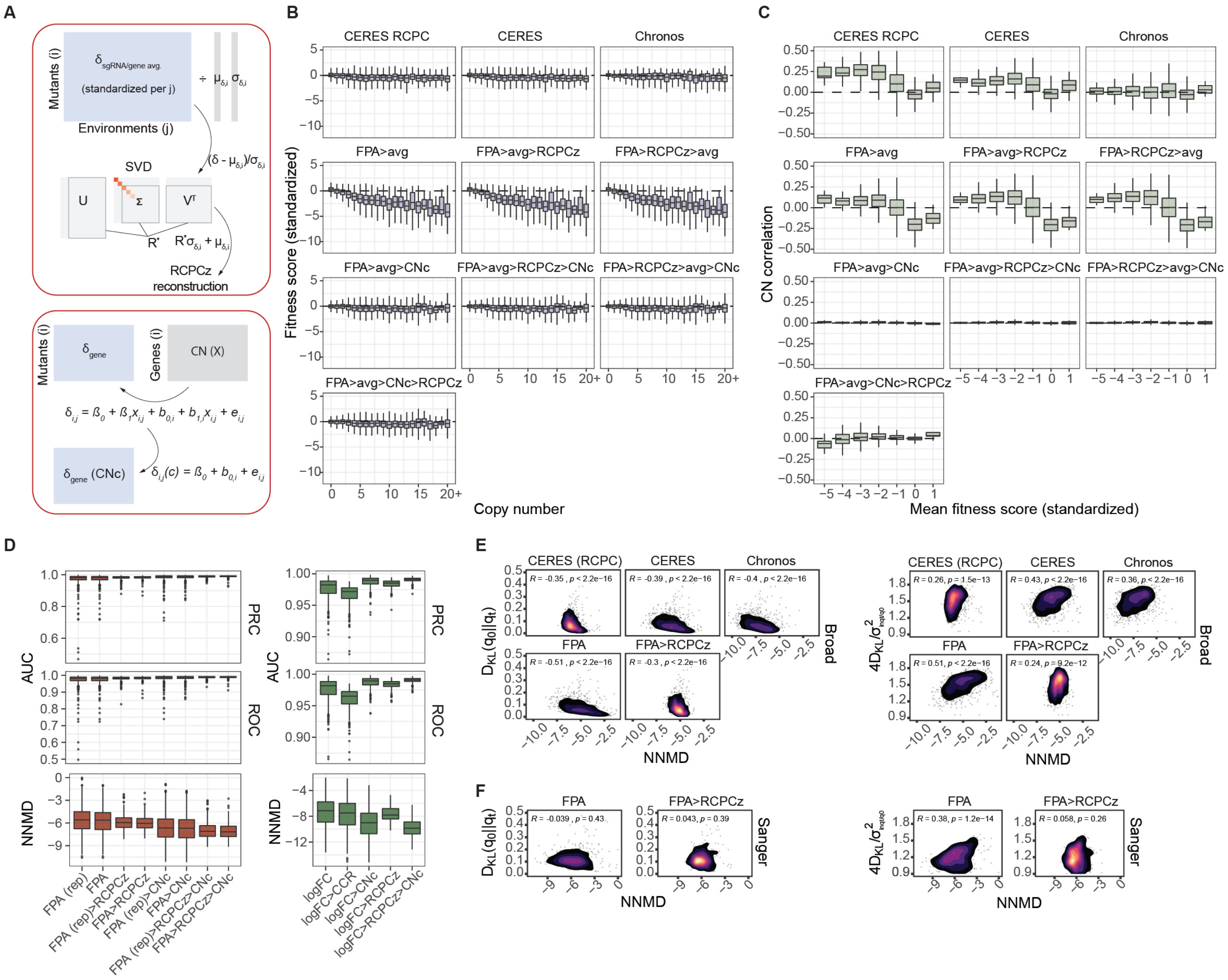


**Fig. S3. Copy number correction and RCPCz for ensemble FPA. A** Overview of the RCPCz procedure (*top*) and the copy number correction procedure (*bottom*). **B** Standardized fitness scores ($\delta(z)$ for FPA) as a function of copy number. **C** Pearson correlation between fitness and copy number per gene, stratified by genes with different mean fitness scores. **D** NNMD, ROC, and PRC AUC for the detection of essential genes comparing fitness computation techniques on the Sanger dataset. *Left panels* show the application of RCPCz and CNc with FPA, and *right panels* show the application of RCPCz and CNc compared to CRISPRCleanR on provided log-fold changes. **E,F** NNMD for detecting essential genes with respect to $D_{KL}$ (*left panels*) or relative divergence (*right panels*) for the Broad (E) and Sanger (F) datasets. $R$ indicates the Spearman correlation.

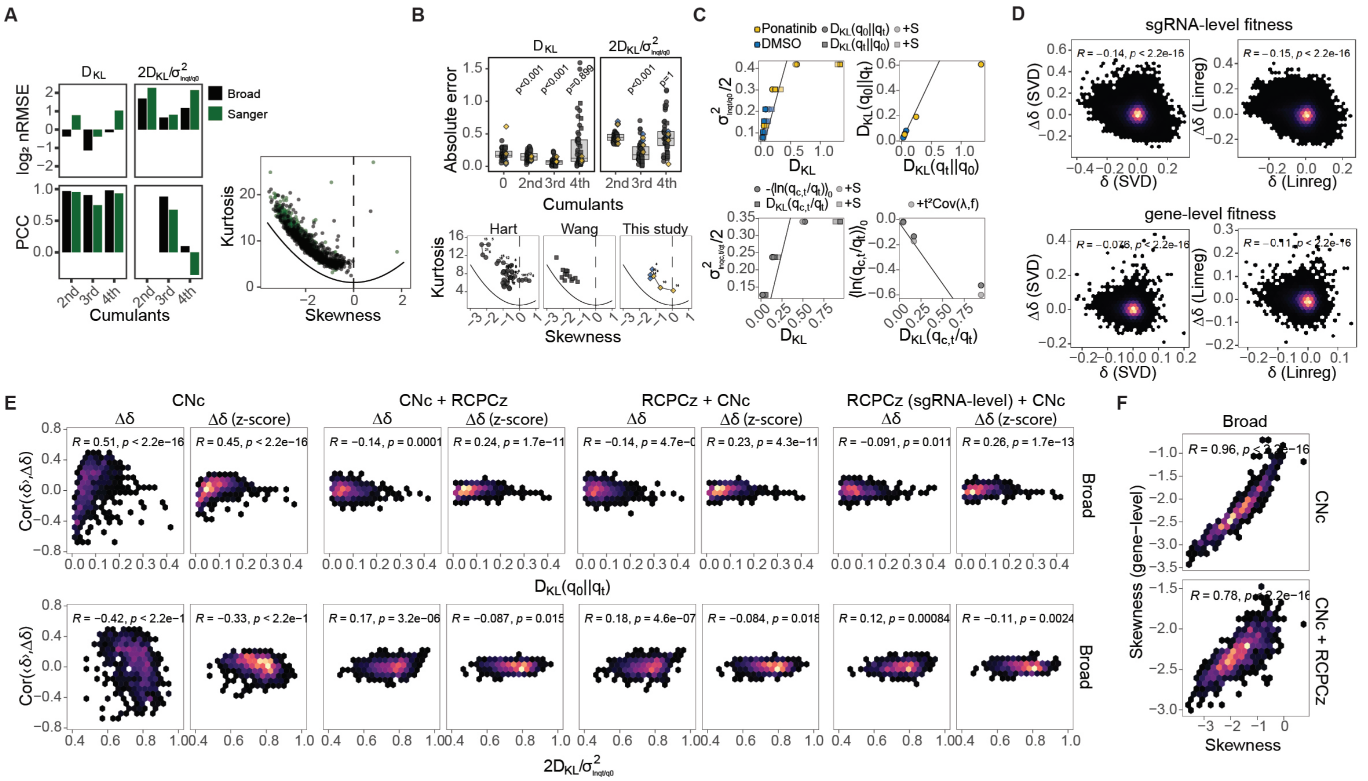

**Fig. S4. Approximation of the $\mathbf{D_{KL}}$ and conditional information. A** Standard deviation-normalized RMSE (*top left*) and Pearson correlation (*bottom left*) for predicted $D_{KL}$ (*left*) or relative divergence (*right*) as a function of increasing cumulants for the Broad and Sanger CRISPR screens. The *right panel* shows the relationship between skewness and kurtosis, with the Cauchy-Schwarz inequality indicated by the *solid line*. **B** Absolute error for predicted $D_{KL}$ (*left*) or relative divergence (*right*) as a function of increasing cumulants (*top panels*) for single CRISPR screens. The *bottom panels* show the relationship between skewness and kurtosis, with the Cauchy-Schwarz inequality indicated by the *solid line*. **C** Relationship between different $D_{KL}$ measures. *Top panels* show prior and posterior $D_{KL}$ with respect to empirical variance (*top left*) and with respect to each other (*top right*), for CRISPR screen samples under selective pressure (Ponatinib) or untreated (DMSO). $+S$ (*top left*) indicates adjustment of $D_{KL}$ values for the third cumulant (skewness) using their respective Taylor coefficients. The *bottom panels* display conditional $D_{KL}$ and $t_0$ expected conditional divergence with respect to empirical variance (*bottom left*) and with respect to each other (*bottom right*). $+S$ (*bottom left*) indicates adjustment for skewness using Taylor coefficients, while $+t^2Cov[\lambda, f]$ (*bottom right*) accounts for covariance between baseline fitness and interaction effects (sgRNA-level). **D** Comparison of baseline fitness and Ponatinib interaction effects for SVD- or regression-estimated values before and after gene averaging; $R$ indicates the Pearson correlation. **E** Cell-wise Pearson correlations between interaction effects and ensemble-wide fitness averages with respect to $D_{KL}$ and relative divergence in the Broad dataset. **F** Gene-level and sgRNA-level skewness in the Broad dataset before and after RCPCz; $R$ indicates the Pearson correlation.

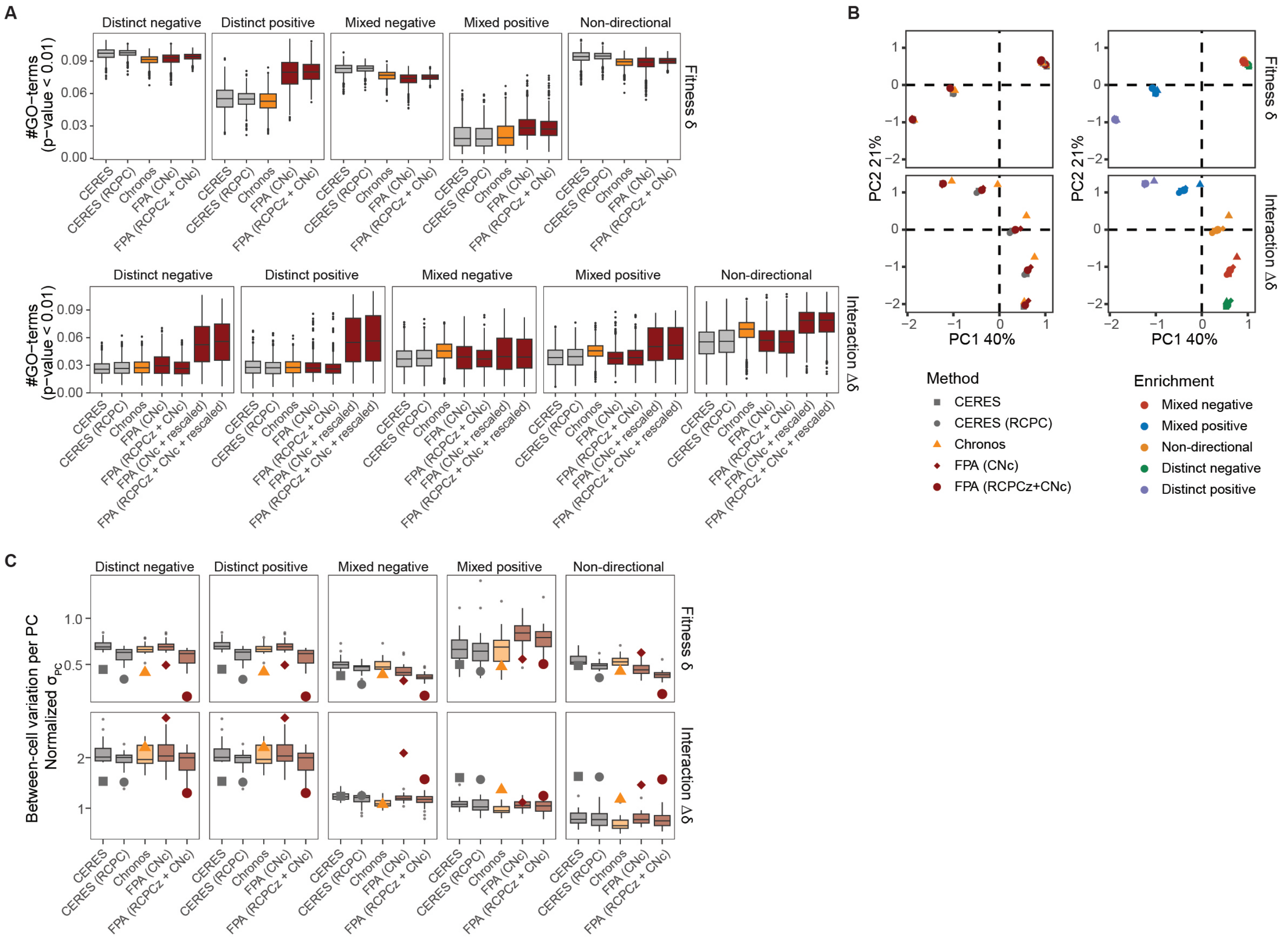


**Fig. S5. Directional GO enrichment benchmarking on the Broad dataset. A** Sensitivity of directional GO enrichment for fitness perturbations (*top panels*) and interaction effects (*bottom panels*) computed using different methods. **B** PCA of enrichment statistics (signed $-\log_{10}(p\text{–}value)$) to compare differences in enrichment results across various fitness computation methods. **C** Variability among cells, assessed by the standard deviation per principal component, for the first components explaining 50% of the enrichment variance. Dots indicate the standard deviations for PC1.

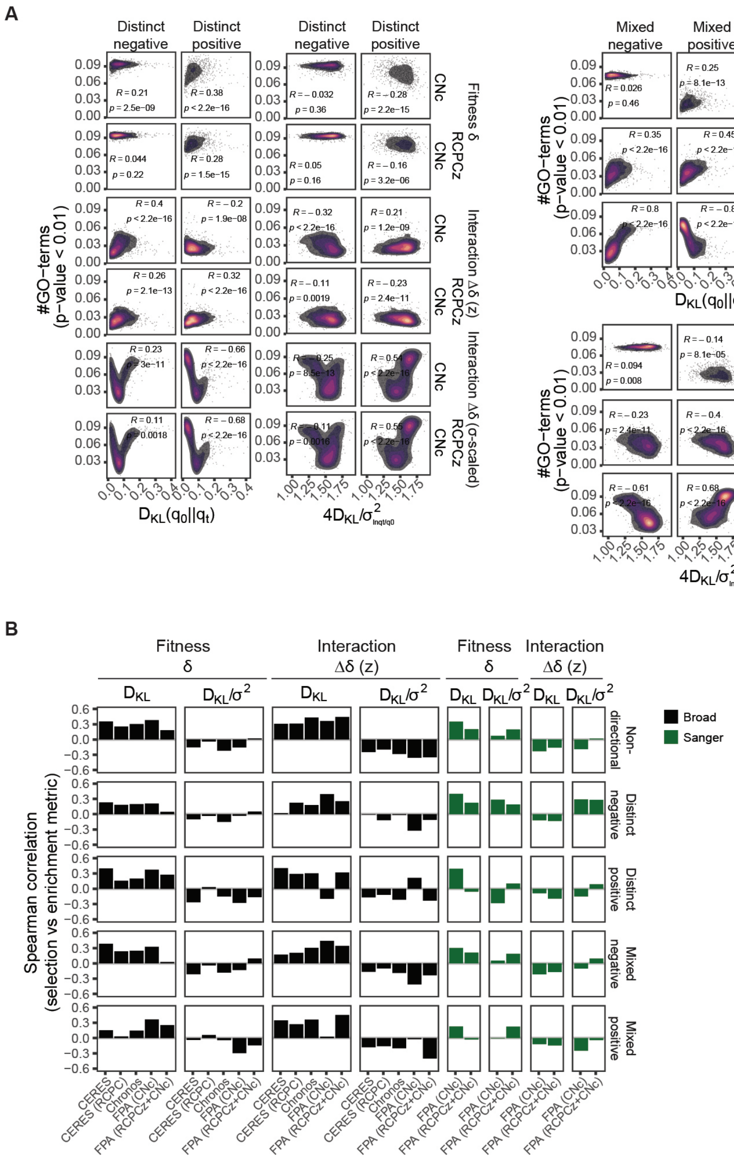


**Fig. S6. Enrichment associations with the $\mathbf{D_{KL}}$.** **A** Sensitivity of fitness GO enrichment with respect to the $D_{KL}$ and relative divergence; $R$ indicates the Spearman correlation. **B** Spearman correlations between GO enrichment sensitivities and the $D_{KL}$ or relative divergence.

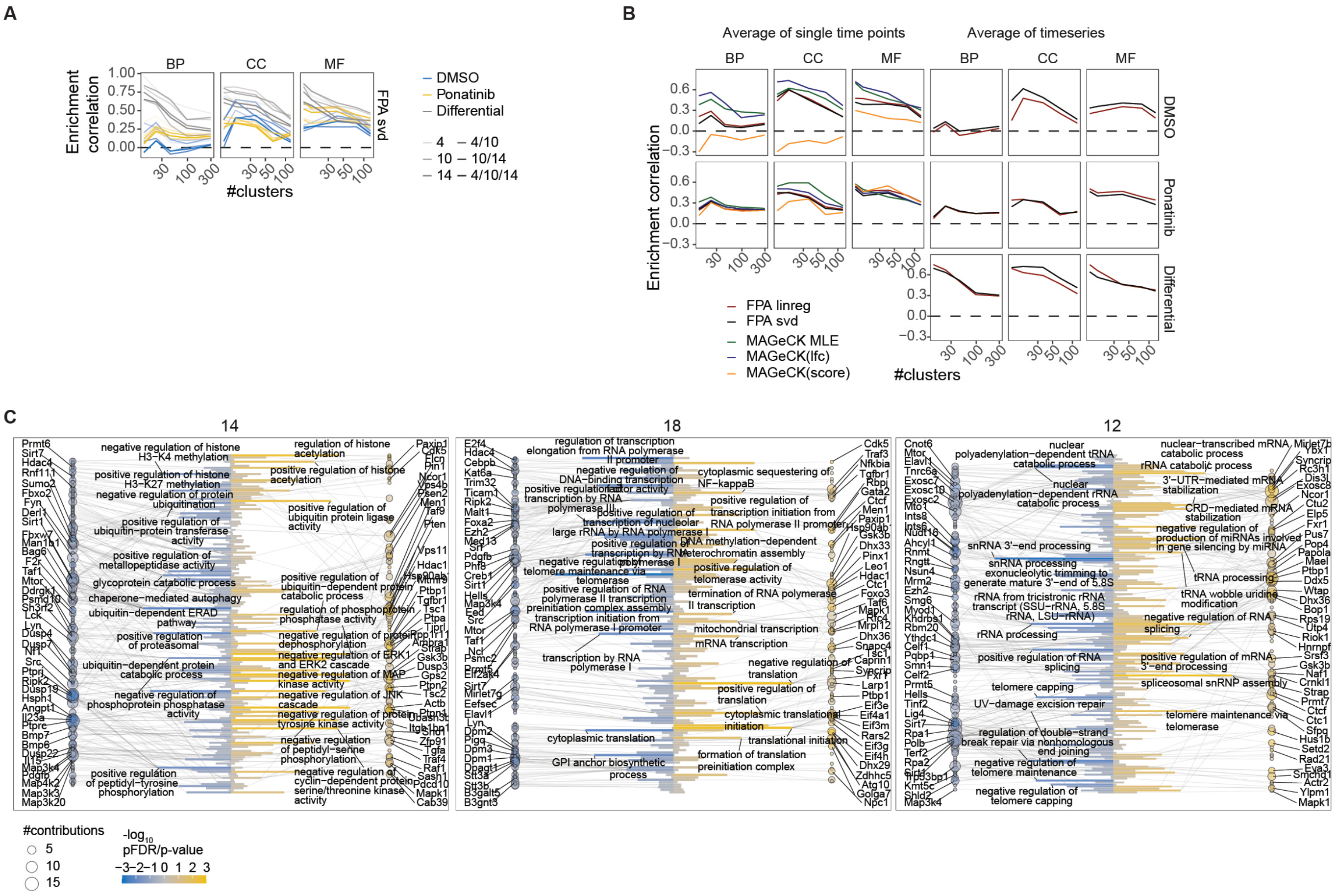


**Fig. S7. Bidirectional enrichment of related functional groups. A** Correlations of directional enrichment across various FPA-estimated fitness perturbations for different clustering depths. **B** Directional enrichment correlations comparing FPA- and MAGeCK-estimated fitness scores. Differential tests for single time points are displayed in Fig. 2D. **C** Directional GO enrichment results for clusters 14, 18, and 12 from Fig. 2C, with contributing genes indicated.

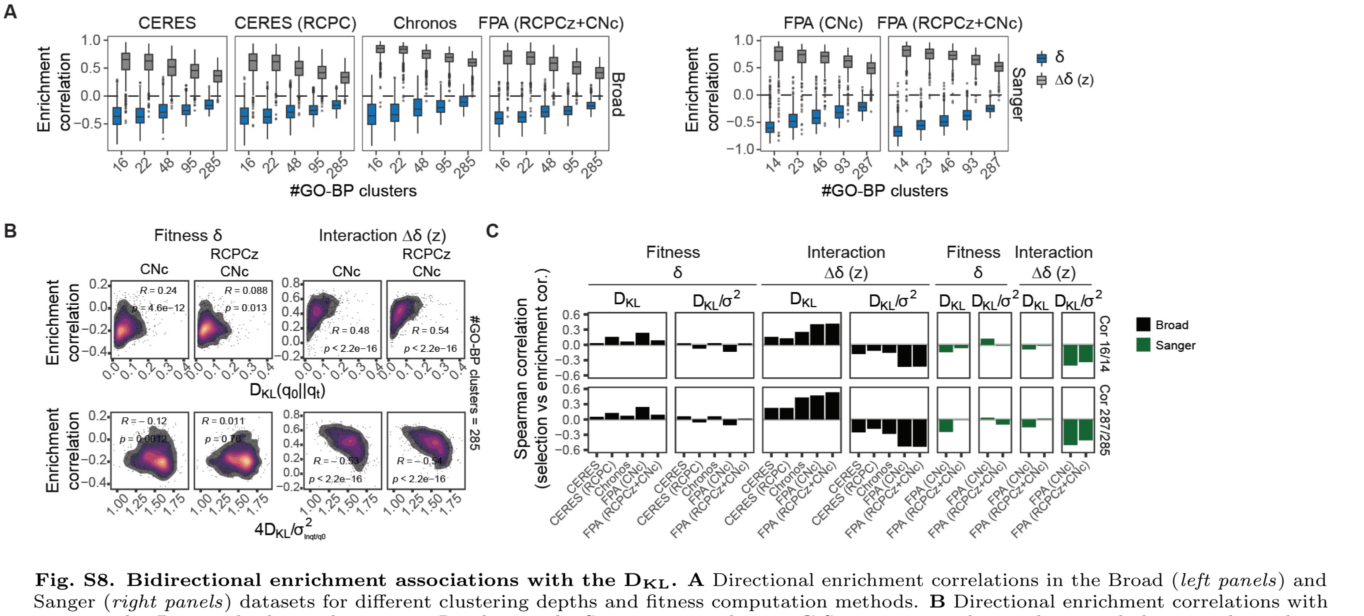


**Fig. S8. Bidirectional enrichment associations with the $D_{KL}$. A** Directional enrichment correlations in the Broad (*left panels*) and Sanger (*right panels*) datasets for different clustering depths and fitness computation methods. **B** Directional enrichment correlations with respect to the $D_{KL}$ and relative divergence; $R$ indicates the Spearman correlation. **C** Spearman correlations between bidirectional enrichment and the $D_{KL}$ or relative divergence.

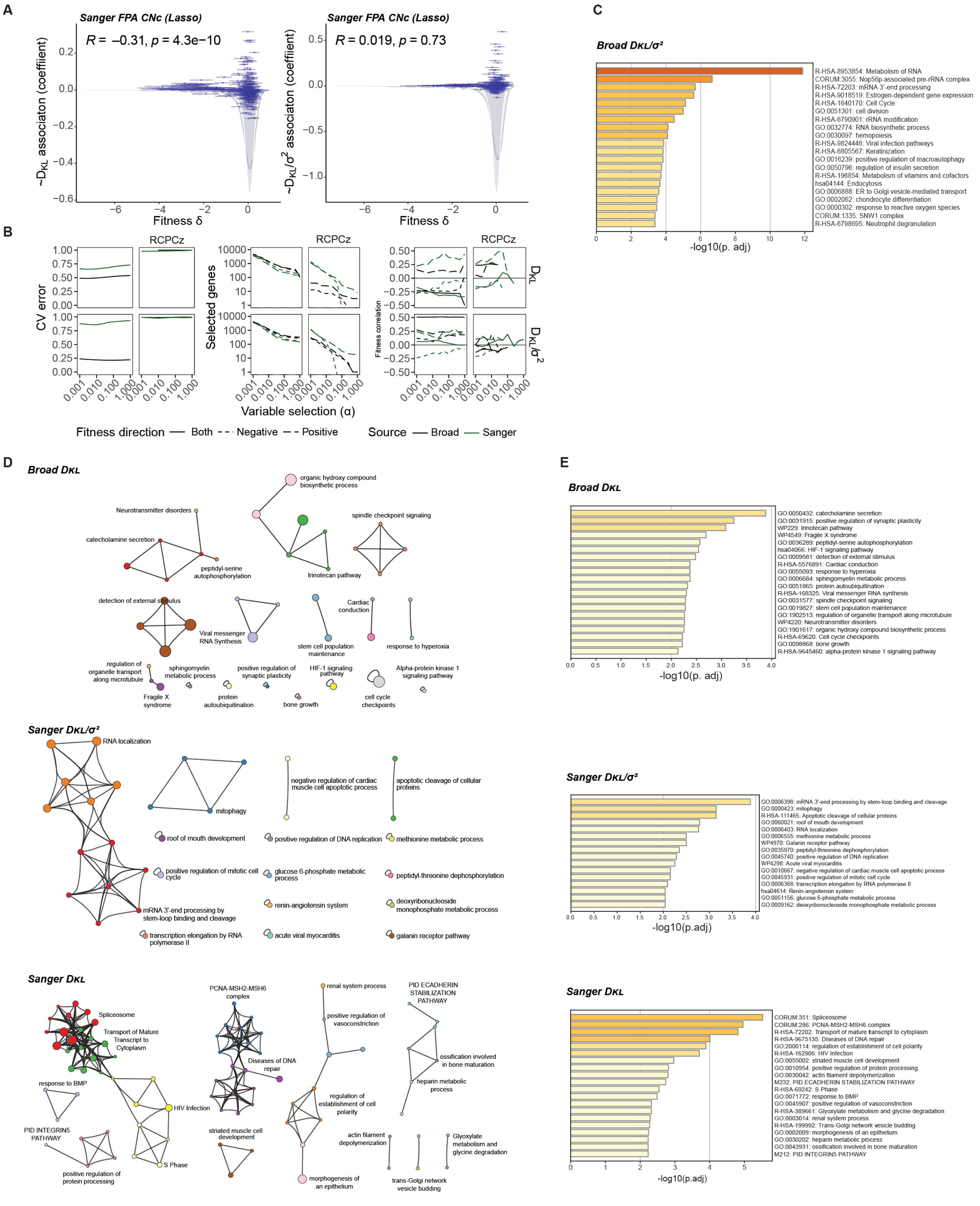


**Fig. S9. Identification of systematic interaction effects driving selection.** **A** Lasso-estimated coefficients for interaction effects with respect to ensemble-averaged fitness perturbations per gene; $R$ indicates the Pearson correlation. The fitness distributions show the selected genes (blue shades) compared to the complete genome (gray lines). **B** Cross-validation (CV) error, number of selected genes, and fitness correlations with regression coefficients for $D_{KL}$ and relative divergence under different variable selection strengths (penalty mixture parameter $\alpha$). **C** Enrichment statistics for clusters shown in Fig. 3C. **D** Enrichment maps for interaction effects predicting $D_{KL}$ or relative divergence. The gene sets were selected using Lasso. **E** Enrichment statistics for clusters shown in panel D.

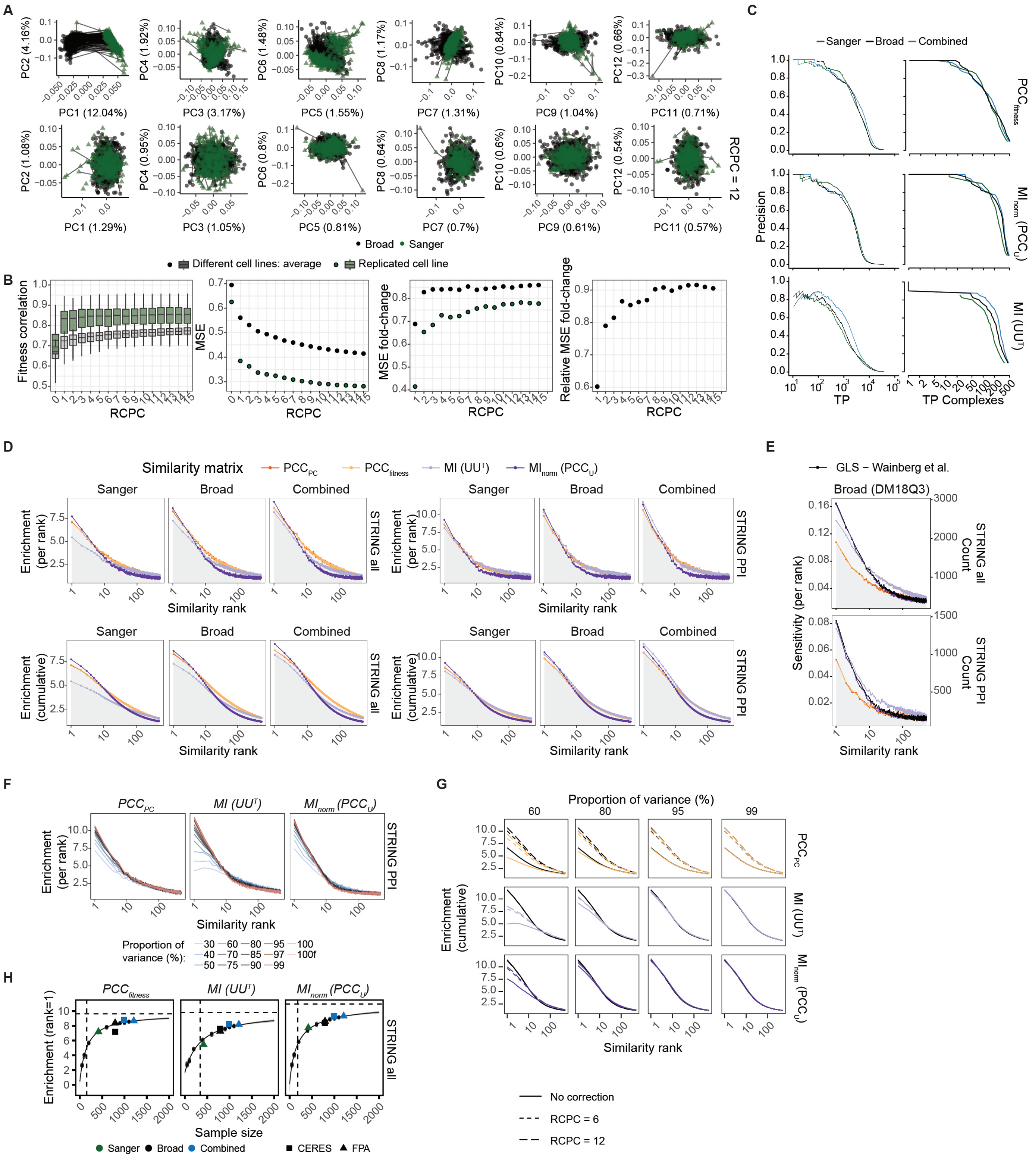

**Fig. S10. Batch correction with RCPCz and enrichment of functional gene interactions. A** Principal component plots for the combined dataset before and after removing 12 confounding principal components. Lines connect replicated cell lines between datasets (*black*) and within the Sanger dataset (*green*). **B** Correlation and mean squared error (MSE) for biological replicates across different conditions as a function of removing confounding principal components. Component-wise relative changes in MSE were used to determine the optimal cutoff ($RCPCz = 12$). **C** Gene-level (*left*) and module-level (*right*) precision-recall curves for retrieval of CORUM complexes using FLEX. **D** Genome-wide enrichment of STRING interactions per individual similarity rank (*top panels*) and cumulative enrichment up to the indicated rank (*bottom panels*) for different functional similarity metrics. Individual datasets were batch corrected with $RCPCz = 6$, while the combined dataset was batch corrected with $RCPCz = 12$. **E** Comparison of genome-wide STRING interaction enrichment between different similarity metrics from this study and GLS from Wainberg et al.[20], using the Broad Institute dataset from DM18Q3. Batch correction was not performed, as indicated by the much lower enrichment scores for $PCC_{fitness}$ and $PCC_{PC}$. **F** Genome-wide STRING PPI enrichment per individual similarity rank for metrics constructed at different perturbation depths, indicated by the proportion of variance explained (%). Results are for the combined dataset with $RCPCz = 12$. **G** Genome-wide cumulative STRING PPI enrichment up to the similarity rank for metrics constructed at different perturbation depths, indicated by the proportion of variance explained (%). Results are shown for the combined dataset without batch correction and with $RCPCz = 6$ or 12. Black lines represent results for full-rank matrices. **H** Genome-wide STRING interaction enrichment per similarity metric as a function of sample size. The curve represents a Hill equation fitted to enrichment scores computed from random subsamples (small dots) of the combined FPA datasets. Horizontal and vertical dashed lines indicate the maximum enrichment and EC50, respectively.

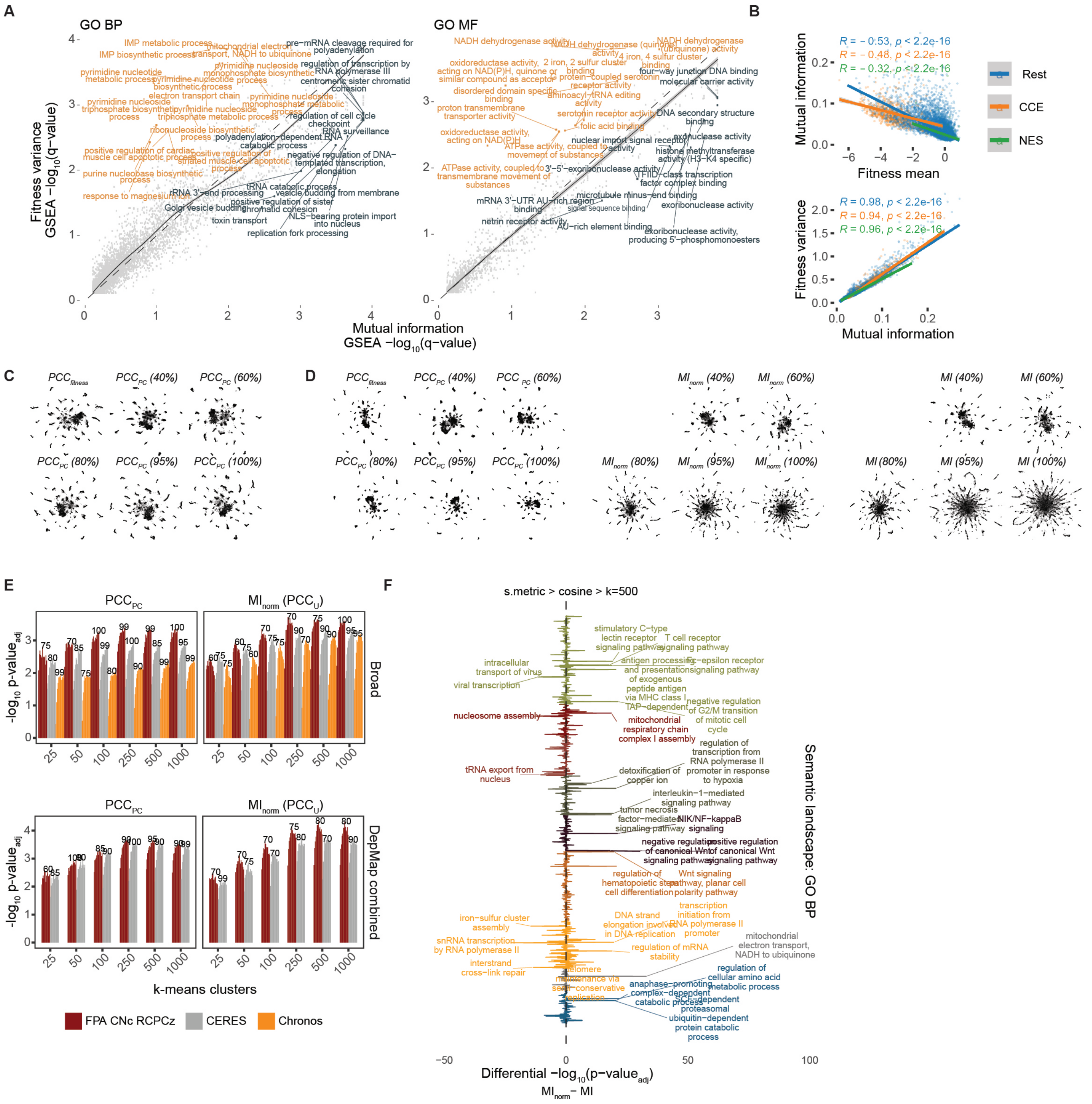


**Fig. S11. GO enrichment and structure of the MI landscape.** **A** GSEA GO-BP and GO-MF enrichment for $I(i;\boldsymbol{\epsilon})$ and $\sigma^2_{\delta_i}$. A LOESS regression line with 95% confidence interval (CI) is shown. **B** Comparison of $I(i;\boldsymbol{\epsilon})$ with $\sigma^2_{\delta_i}$ and $\mu_{\delta_i}$. $R$ indicates the Pearson correlation. **C** UMAP embedding of genome-wide functional landscapes, showing $PCC_{fitness}$ and $PCC_{PC}$ generated by directly embedding $\boldsymbol{\Delta}\delta$ and $\mathbf{g}$ using Pearson correlation as the distance metric. $MI_{PC}$ was embedded with different perturbation ranks corresponding to proportions of variance shown in Fig. 5B. **D** UMAP embedding of genome-wide functional landscapes, showing $PCC_{fitness}$, $PCC_{PC}$, $MI_{norm}$, and $MI$ embedded using cosine similarity as the distance metric. $PCC_{PC}$, $MI_{norm}$, and $MI$ were generated with different perturbation ranks corresponding to proportions of variance shown in Fig. 5B. **E** GO term-wise enrichment summary (average $-\log_{10}(p\text{–}value_{adj})$) for direct UMAP embedding of $PCC_{PC}$ and $MI_{norm}$ based on different perturbation fitness computation methods and k-means clustering configurations. Individual bars represent embeddings with different perturbation ranks corresponding to proportions of variance shown in Fig. 5B, with the configuration showing the highest enrichment labeled. **F** Differential enrichment for GO-BP terms comparing $MI$ and $MI_{norm}$, using a perturbation rank corresponding to 80% of the variance.

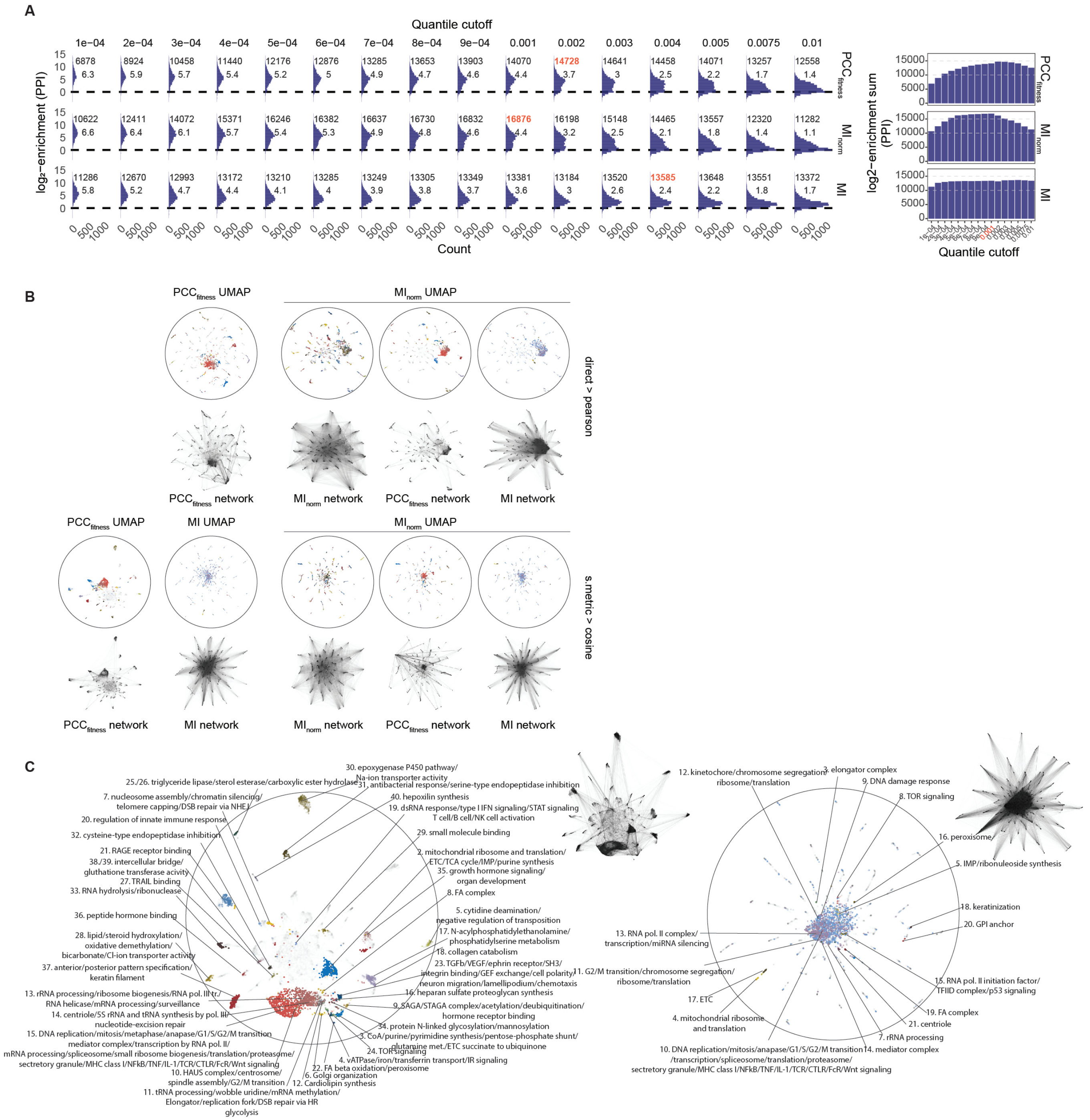


**Fig. S12. GO enrichment across the functional similarity networks. A** Gene-wise enrichment of STRING PPIs for networks generated using different quantile cutoffs. *Left panels* show the distributions, with numbers indicating the integral and median. The highest enrichment is highlighted in red. *Right panels* show the integral per cutoff, with the cutoff used for the subsequent networks highlighted in red. **B** SAFE network domain annotations and network structure for different UMAP embeddings. **C** $PCC_{fitness}$ and $MI$ SAFE network domain annotations and network structure for their respective UMAP embeddings. The similarity matrices were embedded using Pearson correlation as the distance metric.

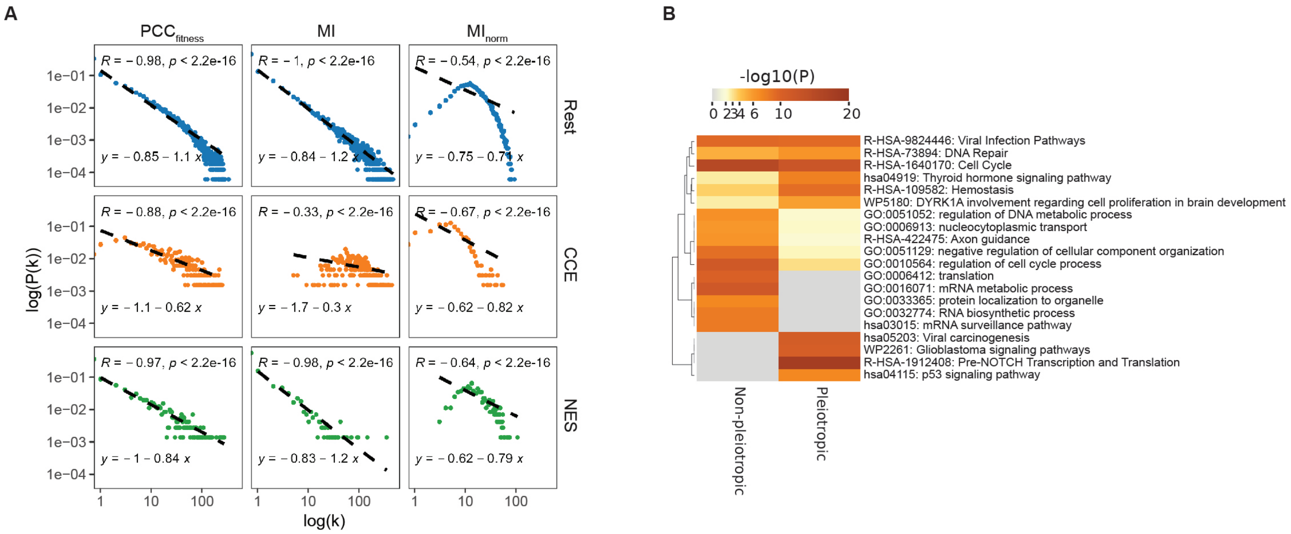


**Fig. S13. Network structure and hub gene characterization. A** Node degree distribution visualized as a log-log plot to assess power-law scaling for each gene essentiality class. $R$ indicates the Pearson correlation, and linear fits (dashed lines) are shown. **B** Metascape enrichment statistics for pleiotropic and non-pleiotropic $MI$ gene hubs.

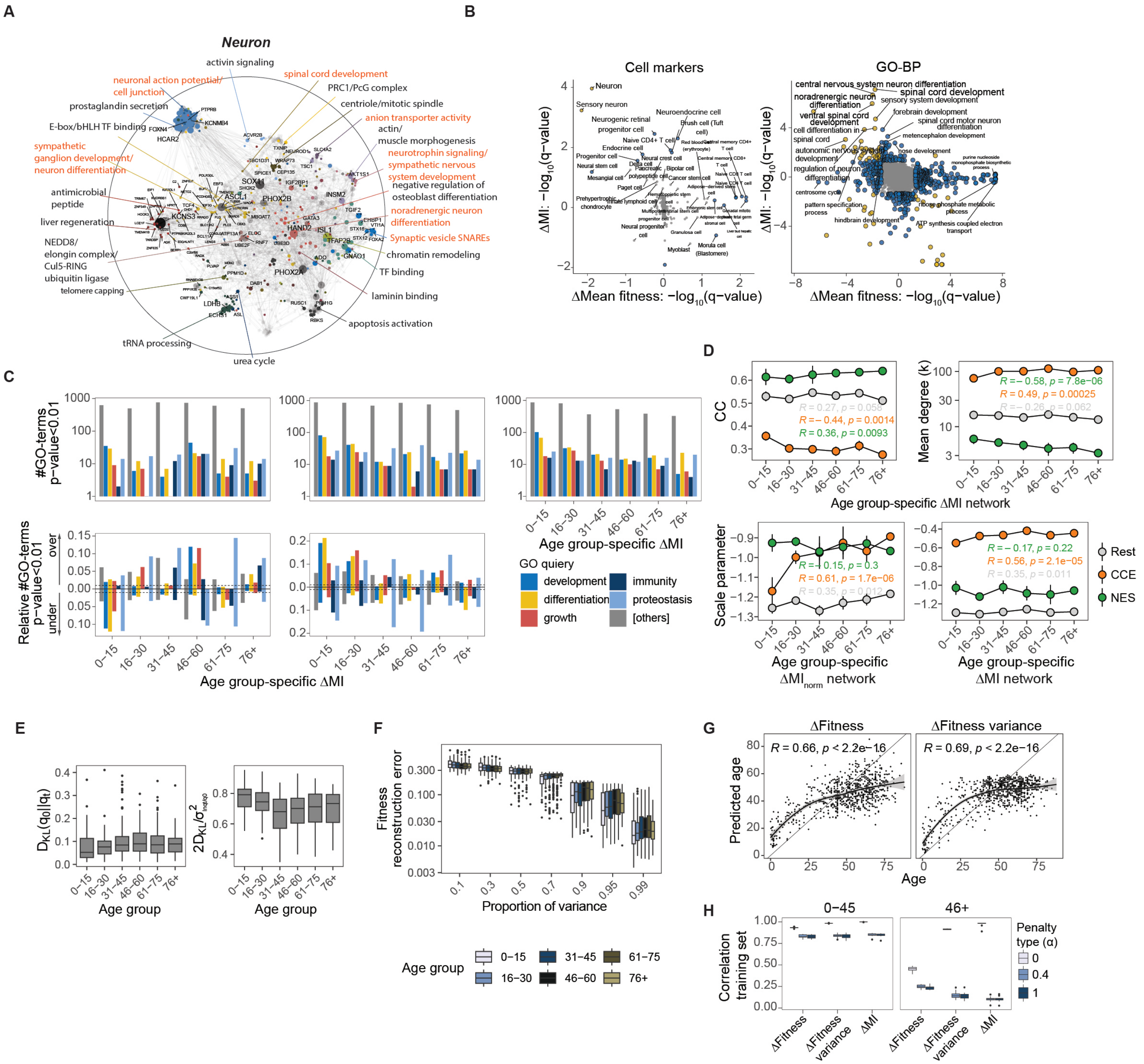


**Fig. S14. Cell-type-specific information captures functional signatures of tissue differentiation, development, and aging. A** $\Delta MI_{norm}$ SAFE network domain annotation for *neurons*. The landscapes represent UMAPs of absolute $\Delta MI_{norm}$ values embedded using Pearson correlation as the distance metric. Labels in red indicate cell-type-specific domains. **B** GSEA cell-type marker enrichment (*left*) and GSEA GO-BP enrichment (*right*) for $\Delta I(i;\epsilon)$ and mean fitness differential for *neurons*. Yellow and blue indicate commonly and uniquely significant terms, respectively. **C** Total count and sensitivity of significant enrichments for indicated GO queries, including $\Delta I(i;\epsilon)$, $\Delta\sigma^2_{\delta_i}$, and $\Delta\mu_{\delta_i}$, stratified by age groups. Directions indicate over- and under-representation. **D** Clustering coefficient (CC) and mean node degree (k) for CCE genes, NES genes, and the remaining genome in age-group-specific $\Delta MI$ networks (*top panels*). Node degree scaling factors are also shown for $\Delta MI$ and $\Delta MI_{norm}$ networks (*bottom panels*). Points and bars represent means and standard deviations from bootstrapped age-group subsets of size $b = 33$, ensuring equal sensitivity across age groups. **E** Screen-wise $D_{KL}$ and relative divergence stratified by sample donor age groups. **F** Fitness reconstruction error stratified by age group for different SVD ranks corresponding to the proportion of variance explained. **G** Lasso-penalized linear regression of age using cell line-specific $\Delta\delta_i$ and $(\Delta\delta_i)^2$ values. $R$ indicates the Pearson correlation comparing predicted age with true donor age. The curve represents a LOESS fit indicating the average prediction per age. **H** Correlation of 10-fold training set predictions of age (log) by penalized linear models trained on different cell-line-specific metrics, stratified by age groups.
